# ON-OFF nanopores for optical control of transmembrane ionic communication

**DOI:** 10.1101/2024.11.23.624940

**Authors:** Xingzao Wang, Aidan Kerckhoffs, Jorin Riexinger, Matthew Cornall, Matthew J. Langton, Hagan Bayley, Yujia Qing

## Abstract

Nanoscale photoswitchable proteins could facilitate precise spatiotemporal control of transmembrane communication and support studies in synthetic biology, neuroscience, and bioelectronics. Through covalent modification of the α-hemolysin protein pore with arylazopyrazole photoswitches, we have produced “photopores” that transition between iontronic resistor and diode modes in response to irradiation at orthogonal wavelengths. In the diode mode, a low-leak OFF-state nanopore exhibits a reversible increase in unitary conductance of more than 20-fold upon irradiation at 365 nm. A rectification ratio of >5 was achieved with photopores in the diode state by either direct or alternating voltage input. Unlike conventional electronic phototransistors with intensity-dependent photoelectric responses, the photopores regulated current output solely based on the wavelength(s) of monochromatic or dual-wavelength irradiation. Dual-wavelength irradiation at various relative intensities allowed graded adjustment of photopore conductance. By using these properties, photonic signals were converted into ionic signals, highlighting the potential applications of photopores as components of smart devices in synthetic biology.

## [Introduction]

Remote control of transmembrane ionic communication can be facilitated by various molecular tools that respond to external stimuli (e.g. light^1–5^, temperature^6^, magnetic^7^ or electrical fields^8^). Among them, optically activated devices fitted with photo-responsive functional groups have garnered attention, which leverage light inputs to achieve spatiotemporal modulation of ionic signaling across membranes^9–11^.

In top-down synthetic biology, ion fluxes across cell membranes can be modulated by nanodevices containing photoswitchable chromophores. For example, natural light-driven ion transporter proteins, such as microbial rhodopsins^12,13^, generate or dissipate electrochemical gradients through chromophores such as all-trans retinal. This enables regulation of the electrical activities of cells in which rhodopsin expression has been introduced, thereby allowing basic biological activities, such as calcium signalling^14,15^ and neuronal firing^16,17^, to be manipulated. These optogenetic tools can also function as integral parts of miniaturized bioelectronic systems with therapeutic potential, for example, by targeting cardiac arrhythmias^18^ or mediating the restoration of vision^19^. Optical controllability can also be introduced to natural ion channels by designing synthetic photoisomerizable allosteric binders that regulate the opening (ON) and closure (OFF) of ligand-gated receptors, such as Shaker K+ channels^5^, ionotropic glutamate receptors^20^, transient receptor potential channels^21^, and L-type Ca^2+^ channels^22^.

In bottom-up synthetic biology, to modulate transmembrane ionic signaling, there is a strong demand for photoresponsive devices featuring modular design, ease of assembly, immediate and reversible ON-OFF responses, sustained ON and complete OFF states, and the potential to respond to additional input stimuli. Fully synthetic systems offer broad scopes of designs and applications, but face challenges in affording efficient, real-time and light-actuated ion movements across membranes. For example, DNA nanopores equipped with a photoisomerizable lid strand were optically activated or deactivated before inserting into lipid bilayers^23^. Synthetic ion transporters with built-in photoswitches moved ions down a transmembrane gradient one at a time^24,25^. Semi-synthetic approaches are also pursued to leverage the natural ON-OFF machinery of protein channels. Chemical engineering of proteins with photoswitches enabled transient light-activated opening of mechanosensitive channels^4^ or timed irreversible formation of protein nanopores^26,27^. New designs are required to achieve prolonged activation and reversible ionic responses to light.

Here, we describe reversible ON-OFF photopores, which exhibit a long open lifetime in the ON state and a low background current in the OFF state. The ON-OFF machinery has been engineered into a protein construct that was not evolved to open and collapse and can be activated in real time when embedded in membranes. This was achieved by covalent modification of α-hemolysin (αHL) monomers with arylazopyrazoles that exhibit almost quantitative photoswitching between E and Z isomers and exhibit high thermal stability in both states. The photopores have been characterized at both the single-molecule and ensemble levels to showcase their ability to permit or restrict transmembrane ionic current in response to light. The pores exhibit behaviors analogous to both electronic diodes (i.e. one-way current flow) and light-tunable resistors (i.e. current modulation) depending on the wavelength of irradiation. The properties of the photopores have been exploited to generate light-to-ionic signal conversion.

## [Results and Discussion]

### Construction of photoreversible nanopores

To construct photopores, we functionalized α-hemolysin (αHL), a heptameric protein nanopore, with photoisomerizable chemical groups on the interior of the channel (**Fig. 1**). The αHL monomers containing a single cysteine residue at position 111, 115, 125 or 129 were separately modified with one of the three types of photoswitches—*o*-fluoroazobenzene (fAzo)^28^, arylazopyrazole (pzH)^29^, or methyl arylazopyrazole (pzMe)^30^—by using thiol-specific chemistry (**Fig. 1a,b, Supplementary Figs. 1 and 2**). Upon assembly into heptamers, the side chains of these residues project into the lumen of the transmembrane β barrel^31^. The modification of αHL monomers with one molecule of either fAzo, pzH, or pzMe was confirmed by liquid chromatography-mass spectrometry (LC-MS) (**Fig. 1c, Supplementary Figs. 3 and 4**). Modified monomers with these photoswitches self-assembled to form homoheptameric pores when introduced into a synthetic lipid bilayer consisting of 1,2-diphytanoyl-sn-glycero-3-phosphocholine (DPhPC) or in the presence of sodium deoxycholate (**Fig. 1a**).

**Fig. 1.**
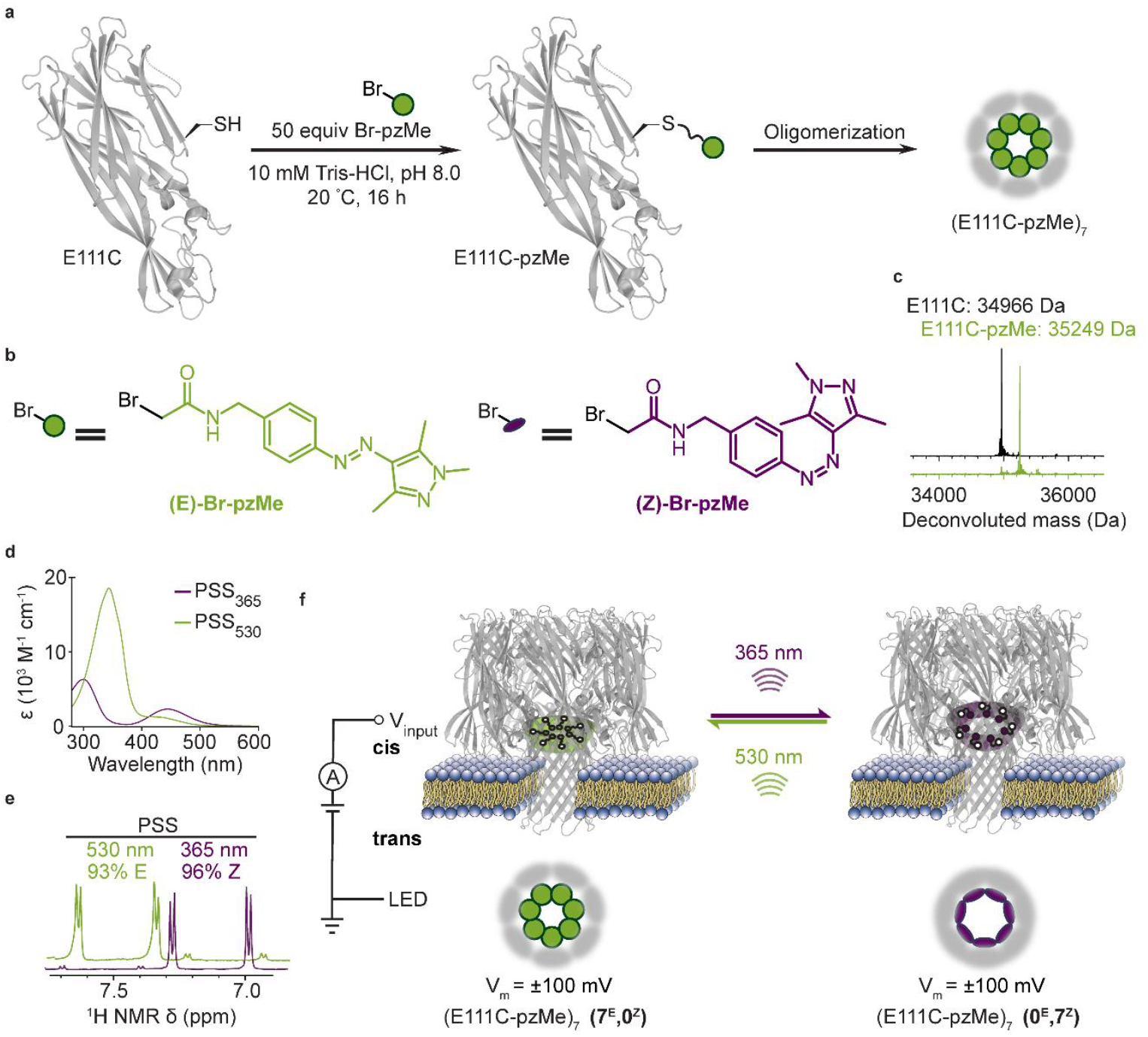
Construction and characterization of the (E111C-pzMe)_7_ photopore. **a**, The α-hemolysin (αHL) monomer containing a cysteine at position 111 was modified with (E)- bromoacetyl arylazopyrazole (Br-pzMe) before oligomerization to form the (E111C-pzMe)_7_ photopore. **b**, The Br-pzMe exists either as the E isomer (green) or the Z isomer (purple). **c**, Deconvoluted mass of E111C and E111C-pzMe monomer confirming quantitative modification of the αHL monomer with pzMe. **d**, UV/Vis spectra of E (green) or Z isomers (purple) of Br-pzMe in DMSO at room temperature. **e**, The photostationary states (PSS) of Br-pzMe examined by the ^1^H NMR chemical shifts of the aromatic protons in DMSO-*d6*. **f**, The modulation of ionic current passing through single or multiple photopores was evaluated by using voltage–clamp electrical recording, in which a transmembrane potential (V_m_) was defined as one in which positive charge moved through the bilayer from the trans side of the pore to the cis. LED light was collimated and projected onto the bilayer from the trans side. Irradiation at 530 nm or 365 nm isomerized arylazopyrazole groups within individual pores, thereby populating photopores in either E or Z states. When all seven arylazopyrazoles adopted the E or Z configuration, such configurations were named as (7^E^,0^Z^) or (0^E^,7^Z^).

**Fig. 2.**
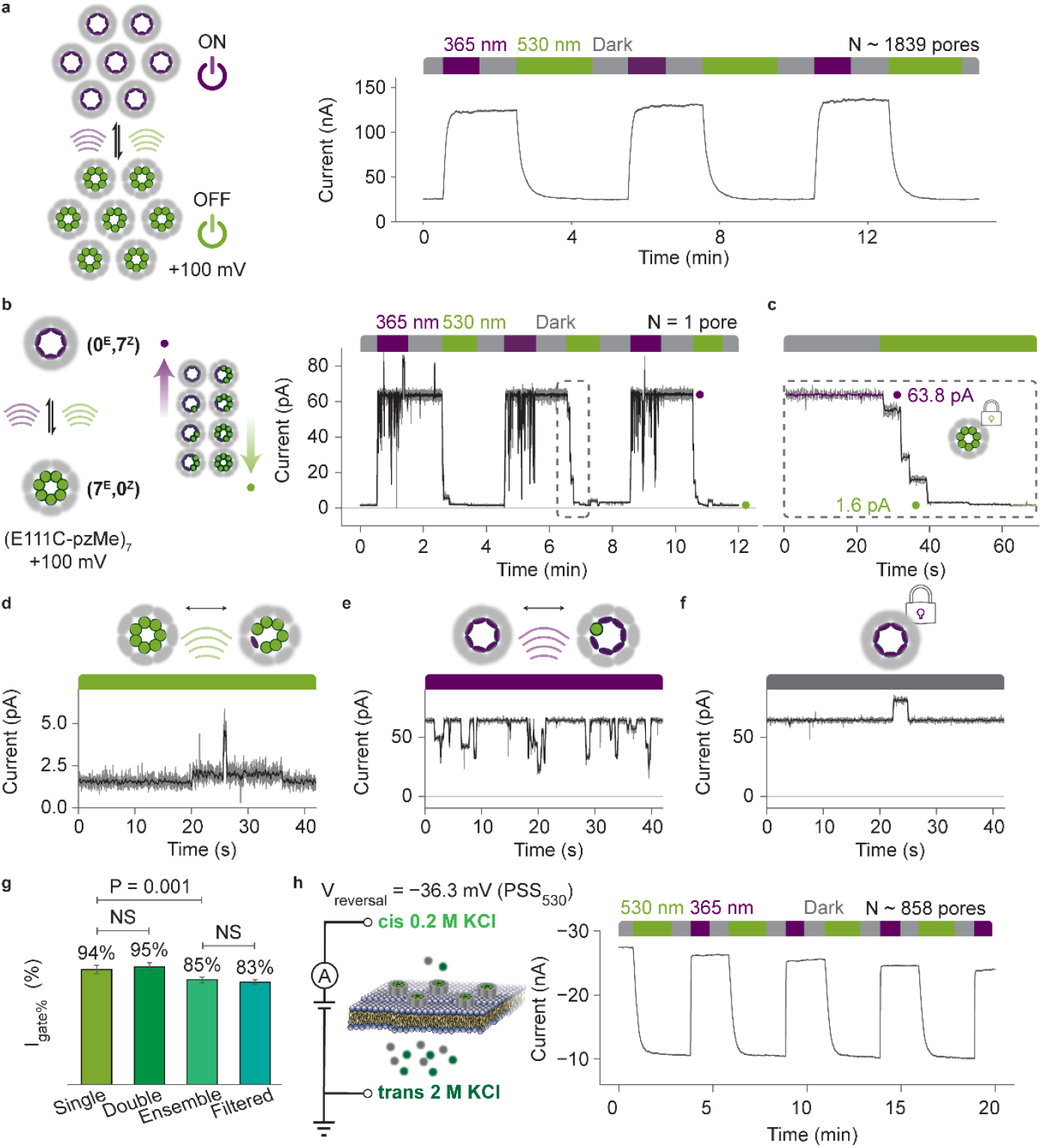
Reversible ON-OFF switching of (E111C-pzMe)_7_ photopore. **a**, Optical control of ionic flow through an ensemble of ∼1839 photopores. Irradiation at 365 nm (purple) converted photopores to an ON state, while irradiation at 530 nm (green) produced an OFF state. The current trace demonstrates ON-OFF switching of the photopore ensemble effected by cycles of UV (365 nm) → dark → green (530 nm) → dark. **b**, Optical control through a single photopore switching between (0^E^,7^Z^) and (7^E^,0^Z^) states. The same light cycle as in **a** was applied. The moving average of the ionic current (grey) is shown in black, which was smoothed with a Savitzky–Golay filter (400 ms window length). During 365 nm irradiation, the photopore switched constantly, causing the black downwards events. By contrast, in the dark, these events stopped. **c**, Zoom-in of the dashed-line box in **b**. Each stepwise decrease in the current was attributed to the isomerization of one or more pzMe photoswitches. **d-e**, Ionic current under continuous irradiation. The current fluctuated due to switching between the (7^E^,0^Z^) and neighboring (6^E^,1^Z^) states at 530 nm in panel **d** or between the (0^E^,7^Z^) and neighboring (1^E^,6^Z^) states at 365 nm in panel **e. f**, Ionic current in the dark. The current remained stable with occasional upward bursts of seconds in duration. Such a dark event was irrelevant to the photoisomerization. **g**, Percent gated current (I_gate%_) in bilayers containing one, two, or an ensemble of pores (n = 4, standard deviations shown as error bars). In-line filters (355-nm filter for 365 nm LED; 532-nm filter for 530 nm LED) were applied to an ensemble of pores with little effect on I_gate%_. **h**, ON-OFF switching of an ensemble of photopores under the electrochemical driving force generated by an asymmetrical KCl concentration across the membrane in the absence of an externally applied transmembrane potential. The current traces were recorded at +100 mV at 25 kHz sampling frequency and filtered using a 5 kHz inline Bessel filter and a 20 Hz digital Bessel filter. Recording conditions: 2 M KCl (**a-f**), 0.2 M KCl (cis)/2 M KCl (trans) (**h**), 10 mM Tris-HCl, 0.1 mM EDTA, pH 8.5, 24 ± 1 °C. The number of pores in ensemble experiments was estimated, assuming each (0^E^,7^Z^) pore contributed +64 pA at +100 mV.

**Fig. 3.**
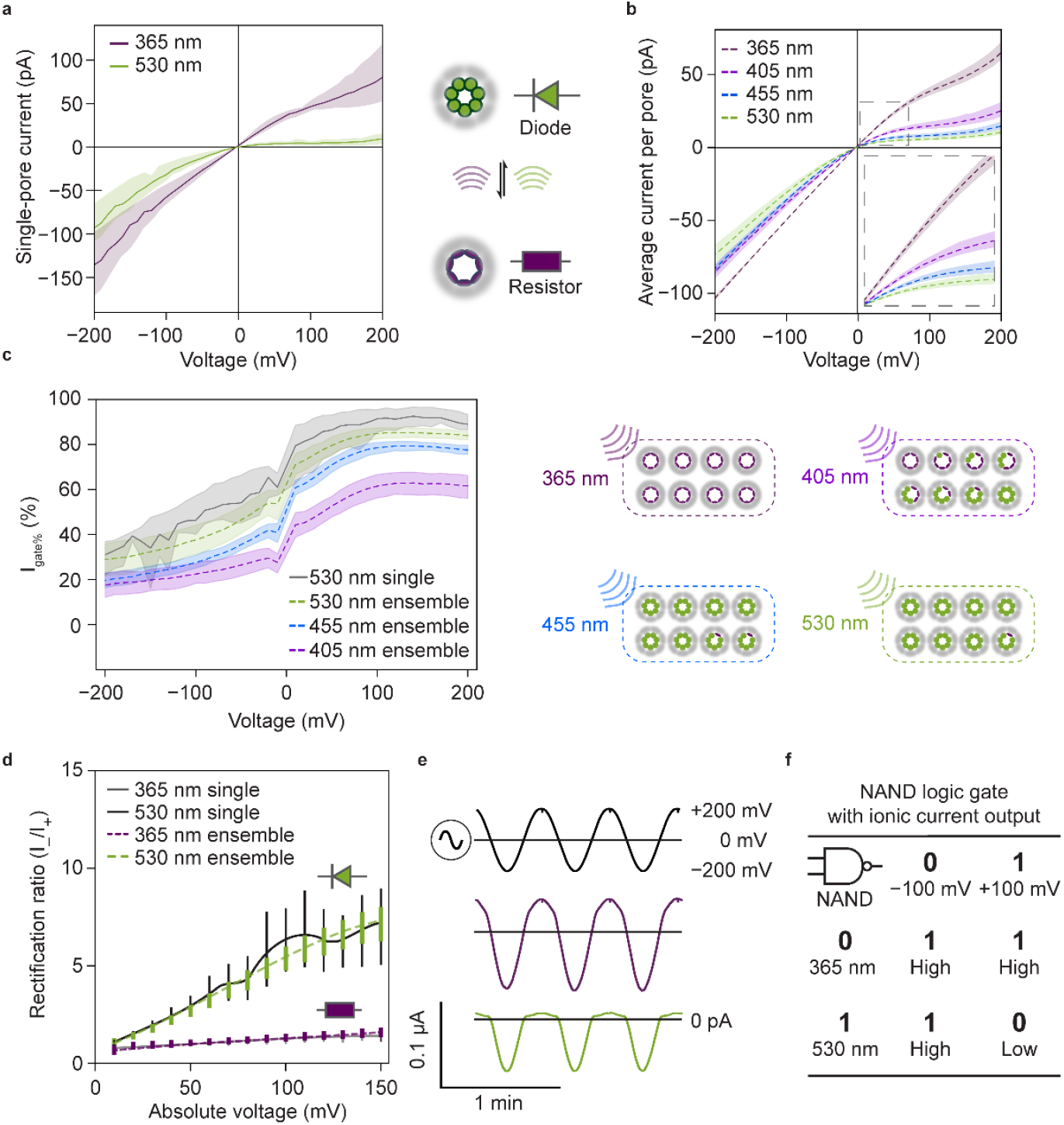
Diode properties of (E111C-pzMe)_7_ photopore. **a**, The current–voltage (I–V) curves of a single (E111C-pzMe)_7_ after 365-nm or 530-nm irradiation reveals photoswitchable resistor and diode behaviors. The 95% confidence interval (N = 3 photopores) is shaded. **b**, The I-V responses of ensembles of (E111C-pzMe)_7_ recorded after irradiation at four wavelengths (365 nm, 405 nm, 455 nm, and 530 nm) and normalized according to the number of pores. Each pore was assumed to contribute +64 pA at +100 mV after 365-nm irradiation. The number of pores in each experiment ranged from 60 to 904 (N = 7 experiments). The 95% confidence intervals are shaded. **c**, Voltage- and wavelength-dependences of the percent gated current [I_gate%_ = (1 − I_OFF_/I_ON_) × 100%], where I_ON_ is the current after 365-nm irradiation and I_OFF_ is the current at the wavelength of interest. The right schematic shows an estimated distribution of intermediate states at PSS of four wavelengths. **d**, Voltage-dependence rectification ratio (I_−_/I_+_) of a single or multiple (E111C-pzMe)_7_ photopores (N = 3 experiments, standard deviations shown as error bars). **e**, Current passing through an ensemble of (E111C-pzMe)_7_ photopores in response to an alternating potential. The curves from the top to bottom are the applied voltage, and the current responses at 365 nm and at 530 nm. **f**, A truth table for the NAND logic achieved using photopores. High and low current levels are achieved by combinations of the input wavelength (λ) and the applied potential. The current I–V curves were recorded with a 5 kHz in-line Bessel filter at 25 kHz sampling frequency, and a 20 Hz digital Bessel filter was used for data analysis. Recording conditions: 2 M KCl, 10 mM Tris-HCl, 0.1 mM EDTA, pH 8.5, 24 ± 1 °C.

**Fig. 4.**
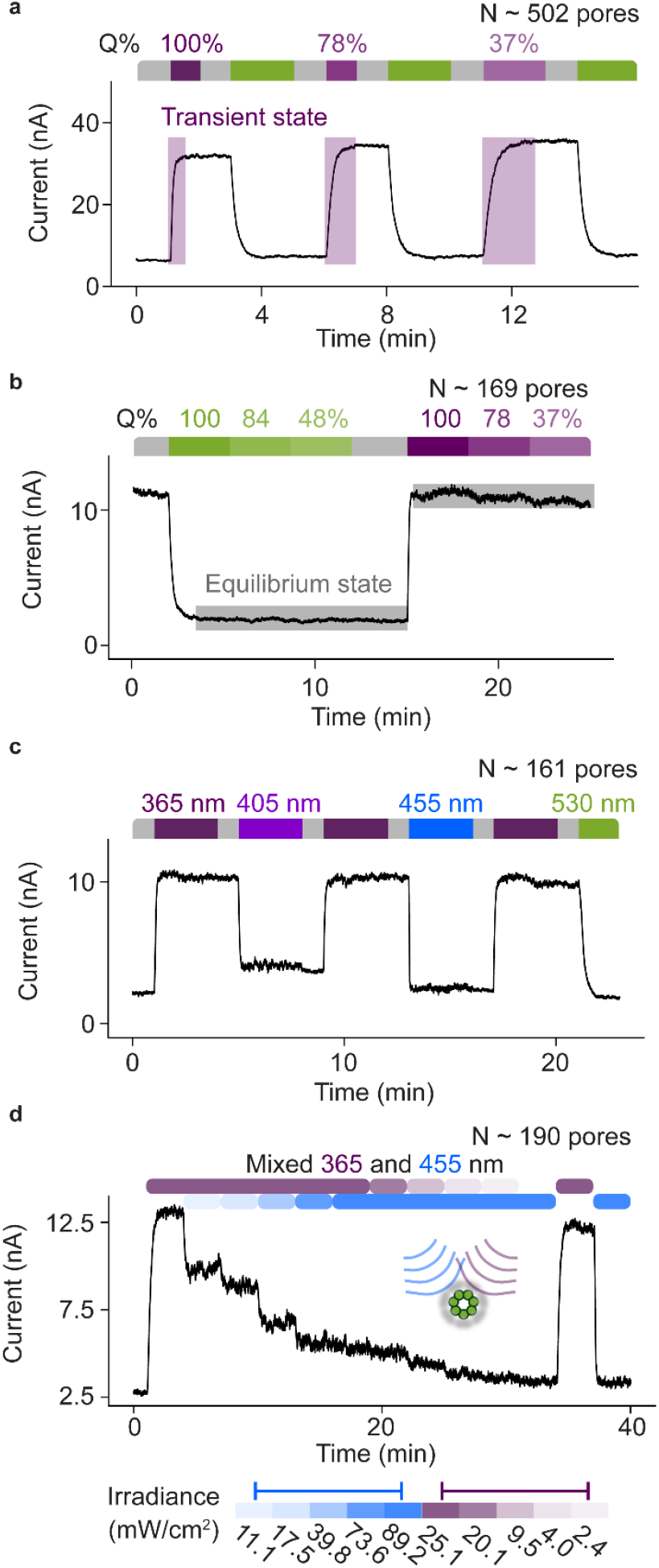
Effects of light intensity and wavelength on the conductance of (E111C-pzMe)_7_ ensembles. **a**, Reduced rate of transition from the low-conductance state (I_OFF_) to the high-conductance state (I_ON_) due to reductions in the intensity of 365 nm irradiation (purple). The percent light intensity relative to the maximum LED intensity (Q%) is shown. The baseline drift is caused by the degradation of Ag/AgCl electrodes when recording for minutes at the nA level. **b**, Insensitivity of output current to light intensity after an equilibrium state is reached. When photopores reached equilibrium states of I_ON_ = 11.4 nA (530 nm, green) or I_OFF_ = 1.8 nA (365 nm, purple) (purple), reductions in the light intensity from the maximum LED output (Q% = 100%) did not cause further changes in the current. **c**, At equilibrium, wavelengths of continuous irradiation result in: an ON state at 365 nm (purple), partially OFF states at 405 nm (violet) and 455 nm (blue) and a fully OFF state at 530 nm (green). **d**, Intermediate current levels can be accessed by polychromic light. Mixed-wavelength irradiation over 3-min intervals with various ratios of 365-nm and 455-nm light allowed fine-control of the PSS ratios with >200 photopores. The current traces were recorded at +100 mV using a 5 kHz in-line Bessel filter at 25 kHz sampling frequency, and a 20 Hz digital Bessel filter was used for data analysis. Recording conditions: 2 M KCl, 10 mM Tris-HCl, 0.1 mM EDTA, pH 8.5, 24 ± 1 °C.

The fAzo, pzH, and pzMe photoswitches were reported to exhibit high E/Z or Z/E isomer ratios at photostationary states (PSSs) as well as thermal stability in the Z state.^28,29^ Determined by ultraviolet/visible (UV/Vis) and proton nuclear magnetic resonance (^1^H NMR) spectroscopy, the bromoacetamide derivative of pzMe (Br-pzMe) demonstrated the highest isomer ratios at PSSs generated by ‘orthogonal’ wavelengths (**Fig. 1d,e**). Upon LED irradiation at 365 nm, Br-pzMe was converted primarily to the Z isomer (PSS_365_ = 96% Z), while when irradiated at 530 nm, Br-pzMe was present predominantly as the E isomer (PSS_530_ = 93% E) (**Fig. 1e and Supplementary Fig. 5**). By contrast, Br-fAzo was 85% in the E form at PSS_405_ and 91% in the Z form at PSS_530_, while Br-pzH was 65% E at PSS_365_ and 85% Z at PSS_530_ (**Supplementary Fig. 5)**. High isomer ratios were expected to produce tight control of ionic communication through modified nanopores.

**Fig. 5.**
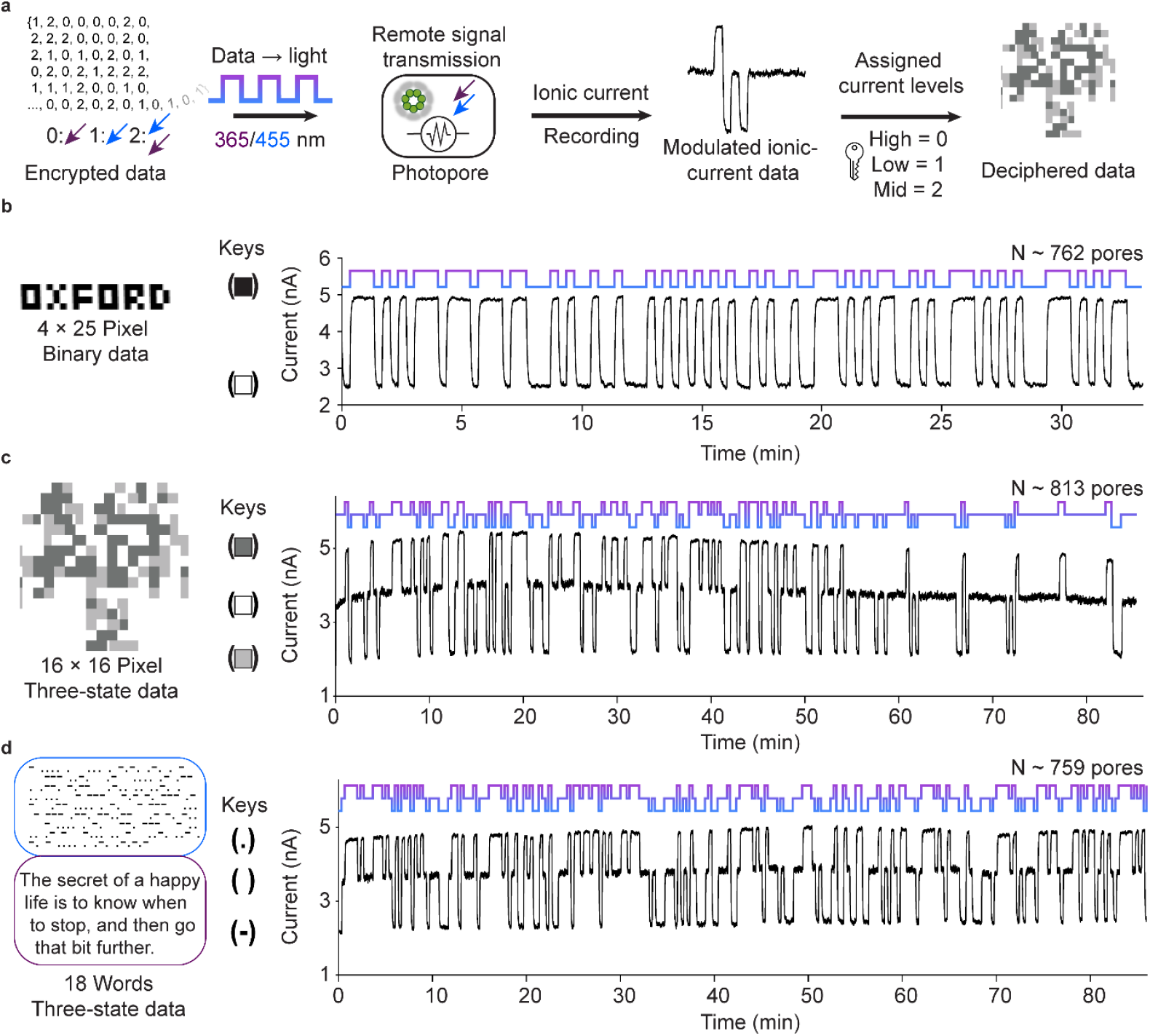
Light-to-current signal conversion. **a**, Overview. Image or text data were encoded as light sequences that produced ionic current responses from (E111C-pzMe)_7_. By using predetermined keys to assign the current levels, the ionic currents were deciphered to reconstruct the original data. **b**, Transmission of a binary pixel pattern with monochromic 365- and 455-nm irradiation. **c**, Transmission of a three-state pixel art in a light sequence. The image of the αHL pore was generated from PDB: 7AHL and converted to three-color pixel art. Each pixel was assigned to 0, 1 or 2 based on the color. The matrix of pixels was flattened to 1D to produce the light sequence. **d**, Transmission of three-state Morse code. The blue box is the Morse code from the deciphered current trace, and the purple box is the translated text. The light sequence was the input at wavelengths of 365 nm, 455 nm or mixed 365/455 nm at a rate of 20 s per bit to generate a current pattern consisting of three levels. The current traces were recorded at +100 mV using a 5 kHz in-line Bessel filter at 25 kHz sampling frequency, and a 20 Hz digital Bessel filter was used for data analysis. Currents from (E111C-pzMe)_7_ ensembles were recorded in 150 mM KCl, 20 mM Tris-HCl, 0.1 mM EDTA, pH 8.5, 24 ± 1 °C.

### Reversible ON-OFF photopores

The photoreversible modulation of ionic current passing through αHL nanopores containing fAzo, pzH, or pzMe photoswitches was first characterized by ensemble electrical recordings in planar lipid bilayers (PLB), i.e. in bilayers containing hundreds or thousands of pores. A bespoke poly(methyl methacrylate) (PMMA) chamber was made, equipped with a collimated fiber-coupled LED, for electrical recording under illumination over a range of intensities from the grounded side (trans) of the bilayer (**Fig. 1f and Supplementary Fig. 6**). To align with the convention of reporting current through αHL pores, we defined a positive current as one in which positive ions moved through the bilayer from the trans side of the pore to the cis (i.e. from the bottom of the barrel towards the vestibule). The transmembrane potential (V_m_) was given as the potential on the trans side relative to the cis side.

Interconversion between a high-current level (I_ON_) and a low-current level (I_OFF_) was recorded at +100 mV with pores equipped with fAzo, pzH, or pzMe under alternating wavelengths (365 nm/530 nm for pzH and pzMe, or 405 nm/530 nm for fAzo) (**Fig. 2a and Supplementary Figs. 1 and 2**). We refer to the lower-current state as the OFF state, in which the ionic flow is partially or almost completely restricted, by comparison with the highest obtainable current level, which is the ON state. In general, the photoreversible nanopores switched to the ON state after irradiation at the shorter wavelength (photoswitches in the Z state), and to the OFF state following irradiation at the longer wavelength (photoswitches in the E state).

Of the homoheptamers modified at position 111 with fAzo, pzH, or pzMe, (E111C-pzMe)_7_ produced the largest percent gated current [I_gate%_ = (1 − I_OFF_/I_ON_) × 100%] of 85% at the ensemble level, whereas the other two photoswitches produced low I_gate%_ values of ∼20% (**Supplementary Fig. 2**). When the pzMe modification was moved to positions 115, 125, or 129, a decrease in I_gate%_ was recorded. Without further investigating the mechanism underlying the position- and photoswitch-dependent efficiency of current gating, we proceeded to characterize the (E111C-pzMe)_7_ photopore and explore its applications.

We investigated the photochemistry underlying the ON-OFF behavior of (E111C-pzMe)_7_ photopores at the single-channel level (**Fig. 2b-f**). Sequential insertions of pre-formed heptamers, pre-irradiated at 365 nm, revealed that the ON-state conductance of individual photopores ranged from 0.12 to 1.27 nS. The average conductance of a pore (0.62 ± 0.30, N = 42 pores) was three times smaller than wild type αHL (1.81 ± 0.17 nS, N = 17 pores) (**Supplementary Fig. 7-8**). Nevertheless, an I_gate_% of 95 ± 2.5% (N = 21) was consistently recorded at the single-channel level across all the (E111C-pzMe)_7_ photopores observed (**Supplementary Fig. 9-10**). Reversible photogating was observed by voltage–clamping individual single photopores and exposing them to ten repeats of UV (365 nm) → dark → green (530 nm) → dark (**Fig. 2b and Supplementary Fig. 11**). Upon UV irradiation, a photopore switched from ON to OFF with a mean response time of 1.8 ± 0.8 s (N = 9 transitions), and the OFF-to-ON transition occurred on average within 31.4 ± 7.7 s (N = 9 transitions) under the green light. The LED irradiance values (365 nm: 36.2 mW/cm^2^; 530 nm: 30.3 mW/cm^2^) were derived from measurements of light intensities in microwatts through a 300 μm-diameter pinhole in place of the bilayer (See **Supplementary Fig. 6**).

The intermediate current levels between I_ON_ and I_OFF_ were likely produced by the stepwise isomerization of individual pzMe molecules (**Fig. 2c**). Based on the number of pzMe switches in either the E or Z state within a photopore, we designated the configuration of an ON-state photopore as (0^E^, 7^Z^), the photopore at the maximum OFF state as (7^E^, 0^Z^), and a partially OFF pore as one ranging from (1^E^, 6^Z^) to (6^E^, 1^Z^) (**Fig. 2b**). Notably, the intermediate current levels recorded with a single photopore during its ON-to-OFF or OFF-to-ON transitions differed from one transition to another (**Supplementary Fig. 11 and 12**). For example, during the stepwise reduction of the current recorded through an (E111C-pzMe)_7_ photopore under 530-nm irradiation, more than 10 distinct current levels were recorded across multiple transitions (**Supplementary Fig. 12**). The complexity was attributed to both the combinatorial possibilities of the E/Z configurations of the seven photoswitches^32^, as well as the conformations of the photoswitches. For example, in the (5^E^, 2^Z^) state, there are three possible arrangements of the two pzMe photoswitches in the Z configuration around the central axis of the pore. Once a single photopore reached a terminal ON or OFF state, the current level exhibited fluctuations under continuous irradiation, while remaining relatively stable in the dark (**Fig. 2d-f**). This was expected as continuous irradiation would drive the reversible isomerization of photoswitches within an (E111C-pzMe)_7_ photopore: for example, switching between (0^E^, 7^Z^) and (1^E^, 6^Z^) states under 365 nm or (7^E^, 0^Z^) and (6^E^, 1^Z^) states under 530 nm. In the dark, the photopore would be locked in a single configuration.

The behavior of the (E111C-pzMe)_7_ photopores at the ensemble level was compared to that at the single-channel level (**Fig. 2g**). We found that the I_gate%_ value was ∼10% lower with multiple pores compared to that of a single pore. This was likely caused by the presence of photopores in partially ON or OFF configurations (e.g., (1^E^,6^Z^) under 365 nm and (6^E^,1^Z^) under 530 nm), as reflected by the PSS_365/530_ value, and by the potential presence of defective pores containing one or more unmodified subunits (**Supplementary Fig. 5**). The incorporation of inline bandpass filters to reduce the LED bandwidths to 10 nm did not improve the I_gate%_ value. By employing an asymmetric KCl concentration across a membrane to establish a physiologically relevant membrane potential (**Supplementary Fig. 13**), the photoswitching behavior was reproduced in the absence of an externally applied transmembrane potential (**Fig. 2h**).

### Switchable iontronic diode properties of (E111C-pzMe)_7_ photopores

The ON-OFF behavior of the (E111C-pzMe)_7_ photopore was further demonstrated in a broader range of applied potential by ramping the voltage from −200 mV to +200 mV at 10 mV per step. We characterized the current-voltage (I–V) behavior of a single (E111C-pzMe)_7_ photopore in the dark after reaching the ON at 365 nm or the OFF state at 530 nm (**Fig. 3a**). After irradiation at 530 nm, the photopore exhibited a diode-like non-linear I–V response; almost no current passed through the pore at positive potentials (0 to +200 mV, V_m_ applied on trans side, **Fig. 1f**), whereas the current demonstrated resistor-like behavior at negative potentials (G = I/V = 0.32 ± 0.11 nS at −100 mV, N = 3 pores) (**Fig. 3a**). After irradiation at 365 nm, a less obvious rectification was observed (G = I/V = 0.46 ± 0.06 nS at +100 mV and 0.58 ± 0.12 nS at −100 mV, N = 3 pores) (**Fig. 3a**). In other words, the photopore functioned as a photoresistor after irradiation at 365 nm and as an iontronic diode after irradiation at 530 nm. We speculate that the voltage-dependent behavior of (E111C-pzMe)_7_ photopores might be attributed to a combination of factors, including the biased orientation of photoswitches by dielectrophoresis and electroosmosis and the potentially different stacking of photoswitches in opposite orientations under nanopore confinement.

The dual-mode feature (i.e., diode and photoresistor) demonstrated with single pores was reproduced with hundreds of pores in ensemble experiments (N = 8 experiments, 100 to 600 pores per experiment) under 365-nm and 530-nm irradiation (**Fig. 3b**). Irradiation at wavelengths of 405 nm and 455 nm was also tested; these wavelengths generated current levels between those produced by 365 nm and 530 nm at all applied potentials (**Fig. 3b**). To give an overview of the voltage-dependent behavior in the ensemble, I_gate%_ was calculated following I_gate%_ = (1 − I_OFF_/I_ON_) × 100%, where the current level after 365-nm irradiation was I_ON_ and that after 405 nm, 455 nm, or 530 nm was I_OFF_ (i.e. I_gate%(405 nm)_ = (1− I_405_/I_365_) × 100%) (**Fig. 3c**). Between −150 mV to +100 mV, I_gate%_ increased with decreasing negative potential and increasing positive potential for all three wavelengths. From +100 mV to +200 mV, I_gate%_ plateaued at ∼85% for 530 nm, ∼80% for 455 nm, and ∼60% for 405 nm (**Fig. 3c**). Under negative potentials, I_gate%_ plateaued at ∼20% for 405 nm and 455 nm, and ∼35% for 530 nm when approaching −200 mV (**Fig. 3c**). The difference in PSS ratios associated with irradiation wavelengths was the primary reason for the wavelength dependence of I_gate%_. Moreover, the voltage dependence should be ascribed to the relative population of the strongly rectifying (7^E^, 0^Z^) state. The rectification ratio of (E111C-pzMe)_7_ (RR = I_−_/I_+_, the ratio of currents recorded under opposite polarities at the same potential) scaled almost linearly with the amplitude of the applied potential up to ±150 mV (**Fig. 3d**), and was 5.5 ± 0.7 at ±100 mV in the diode mode after 530-nm irradiation (**Fig. 3d**).

Nanopore diodes can be used in soft devices as half-wave rectifiers33 to convert alternating current to direct current. We tested our photopore as a switchable iontronic diode to effect a unidirectional current across a synthetic bilayer under an alternating voltage at 0.03 Hz (**Fig. 3e**). After 530-nm irradiation, an ensemble of ∼900 (E111C-pzMe)_7_ photopores functioned as a half-wave rectifier and conducted currents only under a negative potential. After 365-nm irradiation, the current flow replicated the sinusoidal shape of the alternating voltage input, albeit weakly rectified. At higher input frequencies, the capacitive current caused by the bilayer would overwrite the resistive current through the pore, diminishing the rectification. The ionic current output through (E111C-pzMe)_7_ photopores was therefore determined by both the wavelength and the applied potential, resembling the NAND Boolean function. This NAND logic gate leveraged the switchable diode properties of (E111C-pzMe)_7_ photopores, with the irradiation wavelength as input 1 (0 = 365 nm, 1 = 530 nm) and the applied potential as input 2 (0 = −100, 1 = +100 mV). A low current output only occurs at +100 mV after irradiation at 530 nm (**Fig. 3f**).

### Light-to-ionic signal conversion with (E111C-pzMe)_7_ photopores

The rate of photoisomerization of a photoswitch is typically proportional to the light intensity5,34,35. Accordingly, irradiation of more than 600 (E111C-pzMe)_7_ pores at 365 nm at 100%, 78% and 37% relative power of the maximum LED output (**Supplementary Fig. 6**) increased the duration of OFF-ON transition from 5.8 ± 0.5 s to 7.1 ± 0.6 s, and then to 11.4 ± 2.6 s (N = 4 ensembles, **Fig. 4a**). After reaching the equilibrium current level under a given wavelength of irradiation, a reduction in the light intensity led to no change in the current level of an ensemble ionic signal (**Fig. 4b**). This ‘memory’ effect stands in contrast to the transient responses of common electronic components, as well as optogenetics tools.

An ionic signal conducted by multiple (E111C-pzMe)_7_ photopores at an intermediate current level between I_ON_ and I_OFF_ could be achieved by switching off the irradiation immediately after reaching the desired current level during an ON-OFF transition at a given wavelength. However, it was challenging to reproducibly capture a transient current level of interest during rapid ON-OFF transitions. An alternative strategy was devised using polychromic light to produce an equilibrium current level as determined by the PSS of photoswitches (**Fig. 4c,d**). To investigate whether mixed-wavelength irradiation could yield equilibrium current levels unattainable by continuous irradiation at either one of the wavelengths, light from 365-nm and 455-nm LEDs was combined through a fiber-coupler. The graded modulation of ionic signal was demonstrated by varying the relative intensity of the 365-nm and 455-nm irradiation projected onto the multiple (E111C-pzMe)_7_ pores in 3-minute steps. Ionic signal levels between those recorded at PSS_365_ and PSS_455_ were obtained (**Fig. 4d**). This approach allowed for finer control over the graded modulation of ionic current through (E111C-pzMe)_7_ photopores.

Inspired by the ‘memory’ effect and graded modulation of ionic signal, we proposed light-to-ionic signal conversion with an ensemble of iontronic photopores. We first encrypted information-bearing binary data within light sequences that alternated between 365-nm and 455-nm radiation (**Fig. 5a**). Functioning as a light-to-current converter, the (E111C-pzMe)_7_ photopores translated the light sequences into two-level current patterns (**Fig. 5a**), as illustrated by a representative segment of a trace with an input rate of 20 s/bit (**Fig. 5b**). Higher input rates were achieved with increased light intensities, reaching 1.5 s/bit at 365 nm and 1.9 s/bit at 455 nm using the maximum LED outputs. Over a 90-minute experiment, no photobleaching was observed. By associating the ON-state with a black pixel and the OFF state with a white pixel, a pixel art spelling “OXFORD” was reconstructed from a binary light message. Furthermore, adding mixed irradiation at both 365 nm (36.2 mW/cm^2^) and 455 nm (56.3 mW/cm^2^) expanded the alphabet to a three-state system, which was applied to demodulate a pixel art representation of a protein nanopore and a text message written in Morse code (**Fig. 5c,d**). In principle, by using mixed-wavelength irradiation, the (E111C-pzMe)_7_ photopores could output a series of current levels, between the levels at PSS_365_ and PSS_530_, as the basis of complex encoding systems. With our equipment, the current levels fluctuate ± 0.3% in the dark after a 20 Hz digital filter, and the I_gate%_ was 0∼85%. Thus, an upper-level estimate for the number of assignable levels would be ∼300.

## Conclusions

In this work, we constructed protein nanopores with unitary conductance values that changed under remote light control. Among the tested photopores, we achieved quantitative, prolonged, and reversible modulation of ionic currents within (E111C-pzMe)_7_ pores following irradiation at 365 nm (ON, high conductance) and 530 nm (OFF, low conductance). Notably, when an (E111C-pzMe)_7_ pore in the ON state was switched OFF at +100 mV, a >95% reduction in the unitary conductance was observed. Similarly, a current reduction of >85% was observed when multiple (E111C-pzMe)_7_ pores were used in ensemble experiments. This sustained ON state and the almost complete OFF state is highly advantageous for real-time control over transmembrane ionic communication in nanotechnology or synthetic biology applications.

The photopores described here differ from previously described photoregulated nanopores in two aspects. First, the photoswitches controlled the conductance of a fully assembled nanopore reversibly, rather than the process of irreversible nanopore assembly^2^7. Second, the current-voltage characteristics of (E111C-pzMe)_7_ exhibited two switchable states: a resistor-like ON-state, after 365-nm irradiation, and a diode-like OFF-state, after 530-nm irradiation. In the diode mode, there is no delay in the rectification of (E111C-pzMe)_7_ after the reversal of the polarity of the applied potential, in contrast to the previous (non-photoswitchable) arginine-rich αHL diode33.

One impact of photopores in the field of nanoscale iontronics might be in light-to-ionic current signaling. In the case of the (E111C-pzMe)_7_ pore, we have used the ability to produce multiple output current levels, such as the three-state system (blue, mixed, and UV), to demonstrate the ability to reconstruct images or text encoded in light pulses. It follows that by hosting light-modulated nanopores in a 2D array of lipid bilayers, an artificial retina might be created to detect colored images. Our previous work on droplet-hydrogel-bilayer bio-pixels reported a related approach for monochromic 530 nm light^36^.

The photopores might also be incorporated into 3D-printed droplet-based synthetic tissues where the spatiotemporal control of communication between compartments is key to the control of tissue-like behavior^37,38^, There, they could be used, amongst other things, to trigger the release of ions or small molecules^38^, mediate shape changes^37^ or produce motion^39^. We further envisage the development of additional photopores by the conjugation protocol reported here to gain control over additional properties such as selectivity in the transport of small molecules and the ability to mediate the translocation of biopolymers.

## Acknowledgements

This research was supported by a European Research Council Advanced Grant (SYNTISU). J. R. is supported by the Friedrich Naumann Foundation for Freedom, which is supported by the German Federal Ministry of Education and Research. Y.Q. was supported by a Glasstone Research Fellowship and a Fellowship by Examination, Magdalen College, Oxford. A. K. acknowledges the EPSRC Centre for Doctoral Training in Synthesis for Biology and Medicine for a studentship (EP/L015838/1). M.J.L. is a Royal Society University Research Fellow. We thank Howard Lambourne at the Physical and Theoretical Chemistry Laboratory, University of Oxford for fabricating the recording chamber.

## Authors and Affiliations

1. Department of Chemistry, University of Oxford, Oxford, UK Xingzao Wang, Aidan Kerckhoffs, Jorin Riexinger, Matthew Cornall, Matthew J. Langton, Hagan Bayley and Yujia Qing

## Contributions

X. W., M. J. L., H. B. and Y. Q. designed the study. X. W. performed the photopore experiments. A. K. performed the synthesis and the UV/Vis and NMR characterization. J. R. and M. C. contributed to the optimization of experimental conditions. All authors were involved in writing the manuscript.

## Competing interests

X. W., Y. Q., and H. B. have filed a patent describing the photopores and the applications thereof.

