## Supplementary for "ON-OFF nanopores for optical control of transmembrane ionic communication"

### Supplementary Information

### Methods

#### General information

All chemicals were purchased from Merck and used as received unless otherwise stated. Br-pzMe, Br-fAzo and Br-pzH were prepared (see Synthesis and characterization of photoswitches) and used as 3 mM DMSO stock solutions for bioconjugation experiments. Details of synthesis are available in the synthesis section. The fiber-coupled LEDs were purchased from Thorlabs (M405FP1, M455F3 and M530F2). An optical fiber (Ø1000 μm, 0.39 NA) was supplied by Mightex. LC-MS of αHL monomers were recorded on a benchtop QTOF mass spectrometer (Waters Xevo G2-XS) with a ProSwift RP-2H column (ThermoFisher, linear-gradient acetonitrile/water, 0.1% (v/v) TFA, 10 min), and the results were deconvoluted using MassLynx software (Waters). An Arduino UNO microcontroller was programmed to trigger and log the time points of irradiation. Molecular modelling was carried out by using open-source PyMol on GitHub (Schrodinger).

#### Preparation of photopore monomers

#### Site-directed mutagenesis of αHL cysteine mutants

Genes encoding αHL mutants with a single cysteine residue at position E111, T115, T125 or T129 were prepared from the pT7-αHL-$D_8H_6$ plasmid by site-directed mutagenesis (QuikChange II XL, Agilent) with the primers listed in Table S1. The template plasmid have been previously reported[1], where the octaaspartate and hexahistidine tag (-$D_8H_6$) was designed for anion exchange (IEX) and Ni-NTA purification. The polymerase chain reaction was initiated by mixing 5 μL 10 × reaction buffer, 10 ng DNA template, 125 ng forward and reverse primers, 1 μL dNTP mix, 3 μL QuikSolution, 2.5 U PfuUltra HF DNA polymerase and nuclease-free water to make a final volume of 50 μL. After thermal cycling (1 × 95˚C:1 min → 18 × 95˚C: 50 s, 60˚C: 50 s, 68˚C: 5 min → 1 × 68˚C: 7 min), 1 μL DpnI (10 U) was added and the mixture was incubated for 1 h at 37˚C to digest the template DNA. Transformation of *E. coli* was performed by incubating 2 μL of the digested PCR product with 45 μL XL-10 Gold competent cells (Agilent) on ice, followed by 30 s heat shock at 42˚C. Subsequently, a Luria-Bertani (LB)-agar plate containing carbenicillin (50 μg $mL^{-1}$, Merck) was inoculated with the transformed cells. All mutants were verified by DNA sequencing.

#### Expression and purification of αHL cysteine mutants

BL21(DE3)pLysS *E. coli* cells (Agilent) were transformed with the mutated plasmid and inoculated onto LB-agar plates containing antibiotics (carbenicillin, 50 μg $mL^{-1}$; chloramphenicol, 34 μg $mL^{-1}$). A single colony from the plate was picked to inoculate 10 mL LB for the preculture. A 400 mL LB culture containing the same antibiotics was inoculated with 4 mL overnight pre-culture. This expression culture was shaken at 37˚C at 250 rpm for approximately 3 h until the OD reached 0.6, when it was cooled 18˚C before the addition of 2 mL 0.1 M IPTG (Fluorochem) to induce protein expression. The culture was further shaken at 18˚C at 200 rpm overnight. The cells were then harvested by centrifugation in a Beckman J25 centrifuge at 5000 rpm for 20 min at 4˚C and resuspended in 10 mL lysis buffer (50 mM Tris-HCl, pH 8.0, 150 mM NaCl, 10 mM imidazole, 0.1% Triton X-100, 5% glycerol, 2 mM TCEP with an EDTA-free protease-inhibitor tablet (ThermoFisher)). Lysis was then performed by the addition of 250 μL 40 mg/mL lysozyme (ThermoFisher), 2 μL 250 U/μL universal nuclease (ThermoFisher) and 25 μL 2 M $MgCl_2$, and incubation on ice for 1 h. The lysate was sonicated at 40% amplitude for 3 min in a 30 s-ON-30 s-OFF pulse train on ice (VCX 500, Sonics). The supernatant was cleared by centrifugation at 29000 x g for 45 min at 4˚C and transferred to a gravity column containing 1 mL (bed volume) Ni-NTA resin (ThermoFisher). The lysate supernatant and resin mixture was mixed at 4˚C on a platform rotator for 1 h. The column was washed with 2 x 15 mL washing buffer (50 mM Tris-HCl, pH 8.0, 500 mM NaCl, 20 mM imidazole, 2 mM TCEP, 0.1% Triton X-100 and 5% glycerol) and eluted with elution buffer (50 mM Tris-HCl, pH 8.0, 500 mM NaCl, 250 mM imidazole, 2 mM TCEP, 0.1% Triton X-100 and 5% glycerol). The fractions (~10 mL) containing αHL were combined and loaded onto a HiLoad 26/600 Superdex 200pg (Cytiva) SEC column equilibrated with SEC buffer (10 mM Tris-HCl, pH 8.0, 150 mM NaCl, 2 mM TCEP and 5% glycerol) at 4˚C. Fractions containing monomers of αHL cysteine mutants were concentrated to 1 mg/mL and stored at −80˚C as aliquots. The mass of the monomer was verified by LC-MS. On average, 9 mg pure αHL-$D_8H_6$ monomer after SEC purification was obtained from 400 mL culture.

##### Conjugation of photoswitches

Br-pzMe, Br-pzH or Br-fAzo (the photoswitches) were dissolved at 3 mM in dimethyl sulfoxide (DMSO). Purified monomers of the αHL cysteine mutants were buffer-exchanged and diluted to 0.5 mg/mL in low TE buffer (10 mM Tris-HCl, pH 8.0, 0.1 mM EDTA). A photoswitch was added to a monomer solution stepwise over 18 h to a final concentration of 750 µM. The mixture was incubated at 20˚C and cooled to 4˚C before the removal of the excessive photoswitch. The extent of reaction was assessed by LC-MS. The conjugated protein solution was passed through a desalting spin column (ThermoFisher) equilibrated in the low TE buffer to remove excess photoswitch and the eluent was stored at −80˚C.

##### Formation and purification of photopore homoheptamers

The photopore monomer was concentrated with a PES protein concentrator (ThermoFisher) to ~8 mg/mL, and exchanged into low TE buffer (10 mM Tris-HCl, pH 8.0, 0.1 mM EDTA) with a desalting spin column. Heptamerization was performed by adding 10% (w/v) sodium deoxycholate solution to the concentrated protein solution, reaching a final concentration of 6.25 mM. The mixture was incubated at 22˚C for 1 h. To separate oligomerized photopores from the remaining monomers, the mixture was loaded onto a membrane-based strong cation exchange spin column (ThermoFisher), which was equilibrated with 20 mM MOPS, pH 7.5. The sample was eluted with the same buffer containing NaCl (0.2, 0.3, 0.4, 0.6, 2 M) (**Supplementary Fig. 7b**). The homoheptamer eluted at ~0.3 M NaCl, and fractions were combined and frozen at −80˚C. Before electrical recording, the MOPS buffer was exchanged for low TE buffer (10 mM Tris-HCl, pH 8.0, 0.1 mM EDTA).

##### Electrical recording in the planar lipid bilayers

##### Custom recording chamber

Planar lipid bilayer recording experiments were performed in a custom poly(methyl methacrylate) (PMMA) recording chamber. The chamber accommodated an LED light source through an adjustable FiberPort collimator (Thorlabs), allowing accurate alignment and direct irradiation of the aperture in the Teflon film (Ø100 µm). Before recording, the collimator was adjusted to achieve consistent projection of the maximum light intensity onto the aperture through a 3-mm-thick sapphire window (Thorlabs). Fiber-coupled LEDs of various wavelengths (365, 405, 455, 530 nm, Thorlabs) were connected to the collimator through a UV-compatible multimode fiber (0.39 NA, Ø1000 µm, Mightex). In the experiments with mixed wavelengths, a bifurcated fiber bundle (0.39 NA, Ø1000 µm, Thorlabs) was connected to two LEDs. Band-pass filters (Thorlabs) were used for the data presented in **Fig. 2g** to achieve a narrow bandwidth of 10 nm for 355 nm and 4 nm for 532 nm (**Supplementary Fig. 9c**). These filters were loaded onto an integrated in-line reflective fiber optic filter (Thorlabs).

**Single-channel and ensemble experiments**

The Teflon aperture of the recording chamber was pre-treated with hexadecane (~5 µL 1% in pentane). After 10 min, recording buffer (1 mL, 10 mM Tris-HCl pH 8.5, 2 M KCl, 0.1 mM EDTA) was added to both the cis and trans compartments. A lipid bilayer was formed on the aperture by adding 1,2-diphytanoyl-3-sn-phosphatidylcholine (DPhPC, Avanti Polar Lipids) to each compartment. A pair of Ag/AgCl electrodes was placed in each compartment through a salt bridge (3 M KCl in 2% (w/v) agarose). The electrodes were covered with black tape to avoid exposure to light from the LED.

To insert a single pore into the bilayer (**Supplementary Fig. 9a**), homoheptamer (0.1 µL, 0.2 mg/mL) was added to the cis compartment. Once a pore had inserted and showed photoresponse, the buffer was perfused five times to prevent further insertions. To achieve the stepwise insertion of multiple pores, more homoheptamer (10 µL, 0.2 mg/mL) was used (**Supplementary Fig. 8a**).

The characterization of the photopore ensemble was recorded using photopore monomers pre-irradiated at 530 nm for 30 min prior to use to ensure the reproducibility of pore insertion (**Supplementary Fig. 9c**). The monomers (~5 µL 0.2~0.5 mg/mL) were added to the cis compartment while 530 nm LED was switched on. Pore insertion occurred under an alternating applied potential of ±20 mV, and the insertion rate slowed down over 15 min. Excess monomers were removed by perfusion of the cis compartment.

Ionic currents were recorded through Ag/AgCl electrodes connected to a patch clamp amplifier (Axopatch 200B, Axon Instruments). The signal was filtered with an in-line 4-pole low-pass Bessel filter (80dB/decade, 5 kHz). A Digidata 1322A digitizer (Molecular Devices) was used to convert the analogue signal to digital form. The data were analyzed with the pCLAMP 10.3 software suite (Molecular Devices). Electrical traces were plotted with Python (3.8.8), Pyabf (2.3.5), Matplotlib (3.3.4) and Seaborn (0.11.1).

**(E111C-pzMe)$_7$ permeability ratios ($P_{K^+}/P_{Cl^-}$) and electro-osmotic flow**

The current–voltage response of (E111C-pzMe)$_7$ was measured over an applied potential ranging from −200 mV to + 200 mV in 10 mV increments with an asymmetric KCl gradient: 2 M KCl (cis) / 0.2 M KCl (trans) (**Supplementary Fig. 13**). The permeability ratio ($P_{K^+}/P_{Cl^-}$) was determined from the potential at the zero current intercept (the reversal potential, $V_r$) by using the Goldman-Hodgkin-Katz (GHK) equation[2]:

$$\frac{\mathrm{P}_{\mathrm{K}^+}}{\mathrm{P}_{\mathrm{Cl}^-}} = \frac{[\mathrm{Cl}^-]_{\mathrm{trans}} - [\mathrm{Cl}^-]_{\mathrm{cis}} e^{\frac{V_r F}{RT}}}{[\mathrm{K}^+]_{\mathrm{trans}} e^{\frac{V_r F}{RT}} - [\mathrm{K}^+]_{\mathrm{cis}}} \quad (1)$$

where *F* is the Faraday constant; *R* is the ideal gas constant; *T* is 298K.

The magnitude of the electro-osmotic flow ($J_{EOF}$) under symmetrical KCl conditions could then be determined from[3]:

$$J_{EOF} \propto V_m \frac{\frac{\mathrm{P}_{\mathrm{K}^+}}{\mathrm{P}_{\mathrm{Cl}^-}} - 1}{\frac{\mathrm{P}_{\mathrm{K}^+}}{\mathrm{P}_{\mathrm{Cl}^-}} + 1} \quad (2)$$

where $P_{K^+}/P_{Cl^-}$ was 0.15 ± 0.01 (N = 4 repeats).

**Transmembrane signal transmission by light-to-current conversion**

To achieve light-to-current signal conversion, three steps are required. First, the encrypted digital data, as pixel art or text, were converted into a monochromatic or polychromatic 1D light sequence. In the 4 × 25 pixel pattern of the 'OXFORD', the pixel art plotted with Matplotlib was unraveled to a 100 × 1 string of two states (black/white), which were further assigned to '365 nm'/'455 nm' to obtain a light sequence. For example, a list of colors ['black', 'black', 'white', …, 'black'] was converted to a light sequence ['365 nm', '365 nm', '455 nm', …, '365 nm'].

In the αHL pixel art, a snapshot of the PDB:7AHL nanopore model was first converted to a three-color 16 × 16 pixelated image by Adobe Illustrator. The value of each pixel was extracted with Python Imaging Library to generate a 16 × 16 × 3 array, where the 3 was for the RGB color code. This array was unraveled to a 256 × 3 array, and the RGB color code was further replaced by the wavelength. For instance, the RGB color code of white (0,0,0) was assigned to 'mixed (365 + 455) nm'. The assignment of grey and dark grey colors to '455 nm' and '365 nm' lead to a 256 × 1 light sequence. Likewise, the 18-word Morse code sequence (dot/ space/ dash) was converted to the same three wavelengths to generate a light sequence.

In the second step, the 1D light sequence was sent to an Arduino UNO microcontroller to trigger the 365 nm and 455 nm LEDs accordingly. Due to the switching rate of the photopore, each unit of irradiation (a bit) was optimized to last for 20 s to observe the rectangular signals. A higher rate of data transmission might be obtained at increased light intensities. During the lifetime of a bit, the LEDs were coupled through the bifurcated fiber bundle and switched on or off according to the light sequence to produce a high (365 nm), middle (mixed), or low (455 nm) ionic current response through the photophore.

The final step was to process the electrical recording to visualize and analyze the output signal. The electrical recording trace was processed in reverse from the 1 × N current signal to the pixel image or text. A continuous recording trace sampled at 25 kHz was segmented into bits of 20 s (one 20-second bit = 500000 data points of 0.04 ms), and each bit was denoted as 'high', 'middle' or 'low' state based on the mean value compared to manually chosen thresholds. The state values were further replaced by ('black'/'white' for the 'OXFORD' pattern), ('dark grey'/'white'/'grey' for the αHL pixel art) or ('dot'/'space'/'dash' for the Morse code). Finally, the new array was transformed into the proper dimension and displayed with Matplotlib to visualize the image or text.

**Table S1**. Primers used for site-directed mutagenesis.

| Primers | Forward and reverse primer sequences (5′→3′) | Length (nt) | GC % |
|---|---|---|---|
| E111C | GAAATTCGATTGATACAAAATGCTATATGAGTACTTTAACTTATGG<br>CCATAAGTTAAAGTACTCATATAGCATTTTGTATCAATCGAATTTC | 44 | 28 |
| T115C | GATACAAAAGAGTATATGAGTTGCTTAACTTATGGATTCAACGG<br>CCGTTGAATCCATAAGTTAAGCAACTCATATACTCTTTTGTATC | 44 | 34 |
| T125C | GGATTCAACGGTAATGTTTGTGGTGATGATACAGGAAAAATTGGCGGCC<br>GGCCGCCAATTTTTCCTGTATCATCACCACAAACATTACCGTTGAATCC | 49 | 45 |
| T129C | GTTACTGGTGATGATTGTGGAAAAATTGGCGGCC<br>GGCCGCCAATTTTTCCACAATCATCACCAGTAAC | 34 | 50 |

#### Supplementary figures

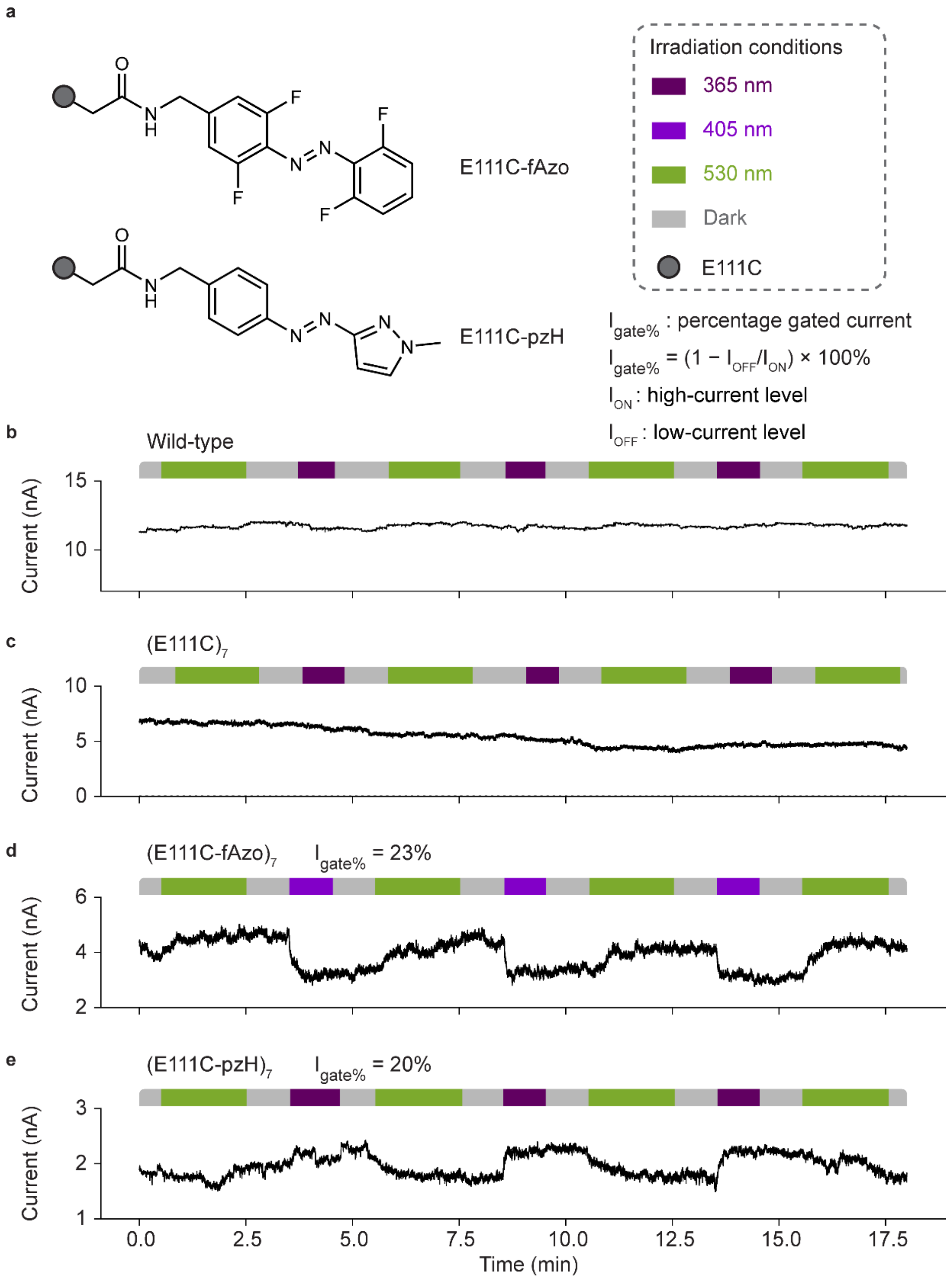

**Fig. 1 | (E111C-fAzo)$_7$ and (E111C-pzH)$_7$ photopores. a,** The fAzo or pzH photoswitches were covalently attached to the cysteine residues at position 111 (E111C) through thioether bond formation. The photoswitches are shown as E isomers. The electrical traces with ensembles of wild-type (WT) α-hemolysin (αHL) (**b**), unmodified (E111C)$_7$ (**c**), (E111C-fAzo)$_7$ (**d**), and (E111C-pzH)$_7$ (**e**) were recorded under alternating light cycles. Wavelengths of irradiation are color-coded: 365 nm, burgundy; 405 nm, purple; 530 nm, green; dark, grey. The WT αHL and (E111C)$_7$ were not photoresponsive. The ensembles of (E111C-fAzo)$_7$ and (E111C-pzH)$_7$ responded to irradiation by E/Z isomerization. The percentage gated current

$I_{gate\%}$ was around ~20%. For (E111C-fAzo)$_7$, irradiation at 405 nm closed the pore, and 530 nm opened the pore. In contrast, for (E111C-pzH)$_7$, irradiation at 530 nm closed the pore, and irradiation at 365 nm opened the pore. The current traces were recorded at +100 mV using a 5 kHz in-line Bessel filter at 25 kHz sampling frequency, and a 20 Hz digital Bessel filter was used for data analysis. Recording conditions: 2 M KCl, 10 mM Tris-HCl, 0.1 mM EDTA, pH 8.5, 24 ± 1 °C.

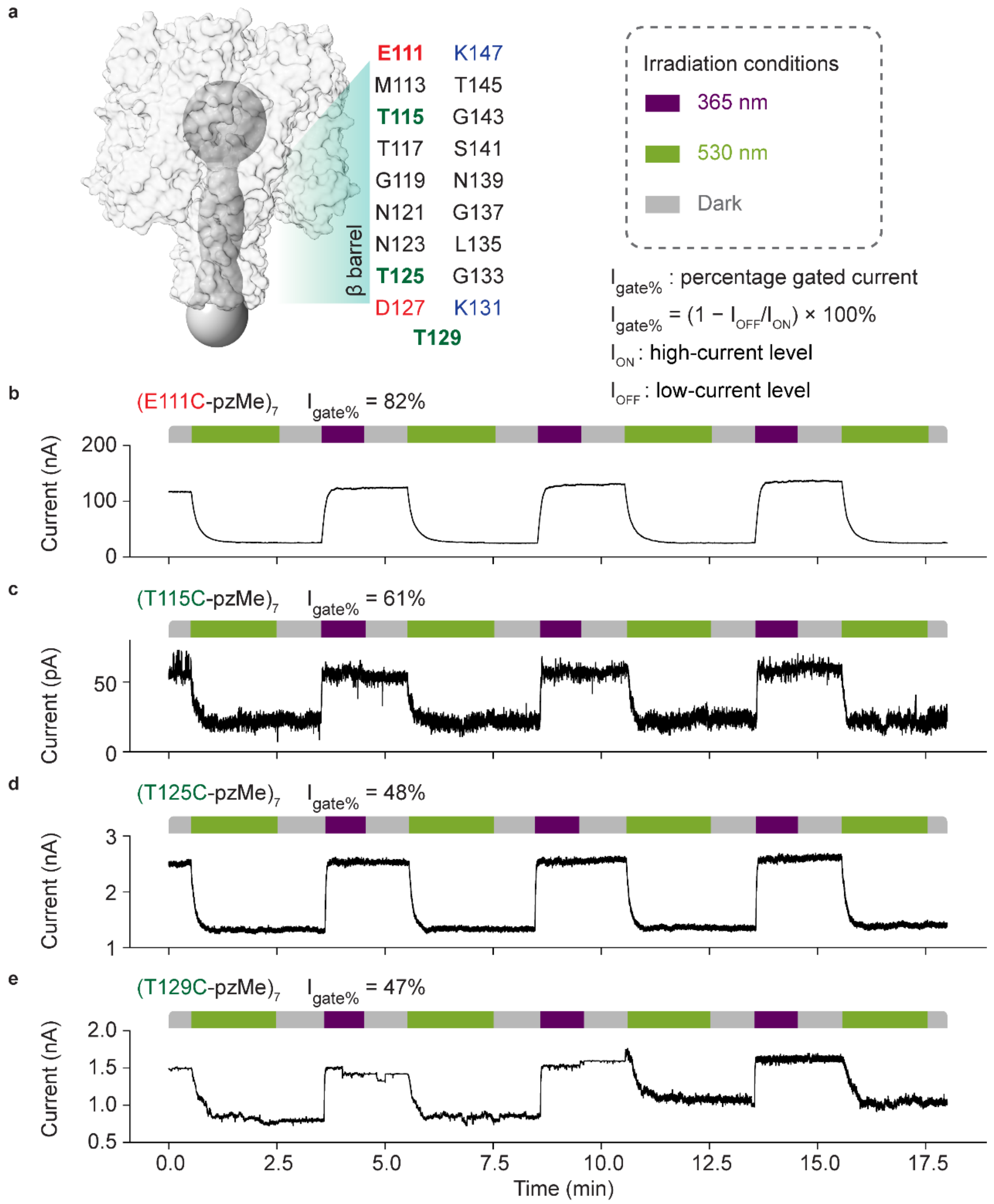

**Fig. 2 | Photopores with pzMe modification in the αHL β barrel. a,** Representation of the αHL pore and residues in the β barrel. The grey semi-transparent structure of the αHL heptamer was generated from PDB: 7AHL with PyMOL. The grey solid dumbbell model showing the protein channel volume was retrieved from ChannelsDB[4]. Electrical traces from ensembles of $(E111C\text{-}pzMe)_7$ (**b**), $(T115C\text{-}pzMe)_7$ (**c**), $(T125C\text{-}pzMe)_7$ (**d**) and $(T129C\text{-}pzMe)_7$ (**e**) were recorded under alternating light cycles. Wavelengths of irradiation are color-coded: 365 nm, burgundy; 530 nm, green; dark, grey. The $(E111C\text{-}pzMe)_7$ showed the largest percentage gated current ($I_{gate\%}$) at 82% in the illustrated trace. All pzMe photopores showed the following correlation—Z, ON; E, OFF. The current traces were recorded at +100 mV using a 5 kHz in-line Bessel filter at 25 kHz sampling frequency, and a 20 Hz digital Bessel filter was used for data analysis. Recording conditions: 2 M KCl, 10 mM Tris-HCl, 0.1 mM EDTA, pH 8.5, 24 ± 1 °C.

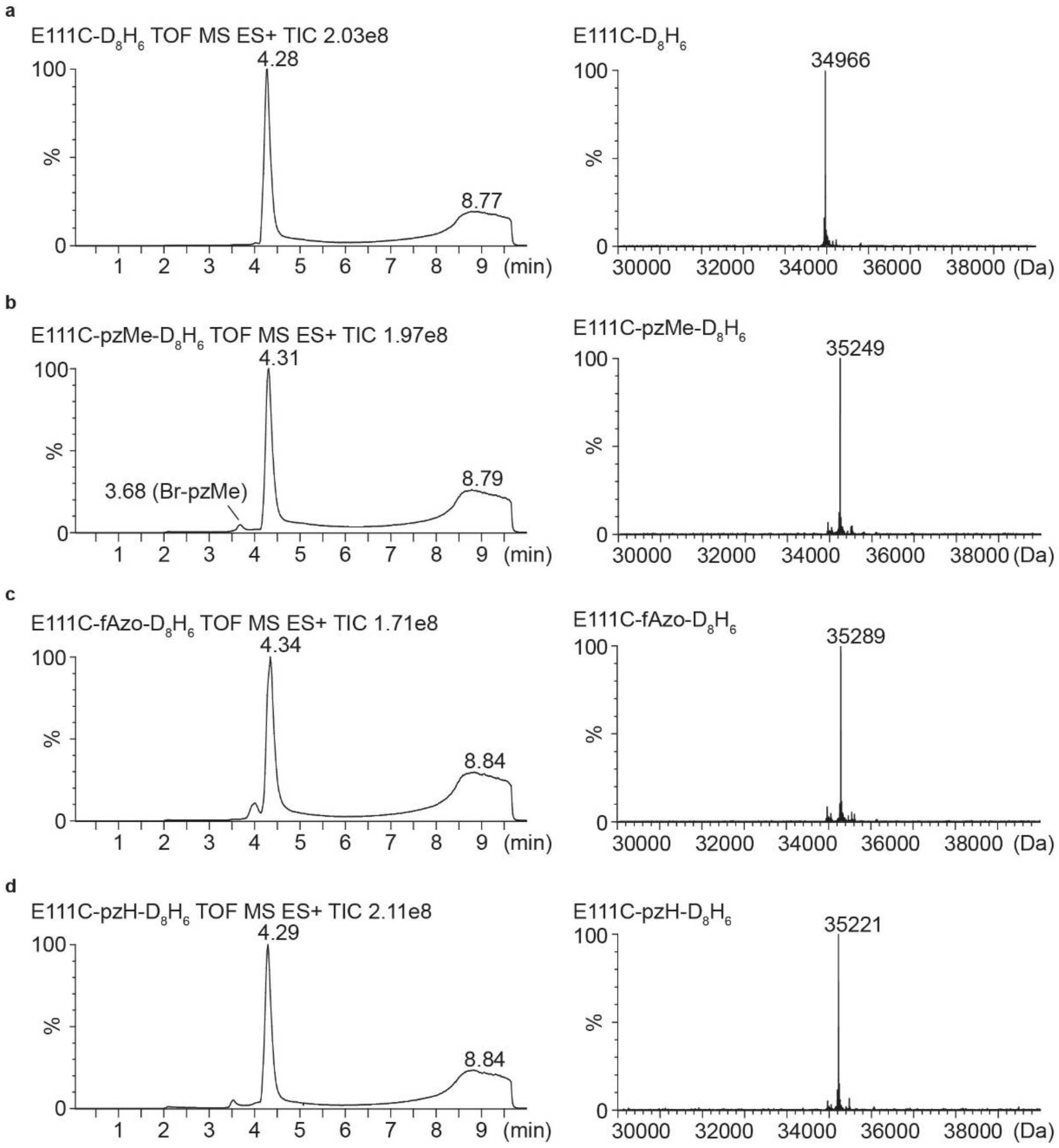

**Fig. 3 | LC–MS characterization of E111C monomers before and after chemical modification.** The left panels are total ion current chromatograms. The right panels are deconvoluted ESI–MS spectra in the range of 30-40 kDa. **a,** E111C-$D_8H_6$ monomer: mass = 34966 (calculated) and 34966 (observed). **b,** E111C-pzMe-$D_8H_6$ monomer: mass = 35249 (calculated) and 35249 (observed). **c,** E111C-fAzo-$D_8H_6$ monomer: mass = 35289 (calculated) and 35289 (observed). **d,** E111C-pzH-$D_8H_6$ monomer: mass = 35221 (calculated) and 35221 (observed).

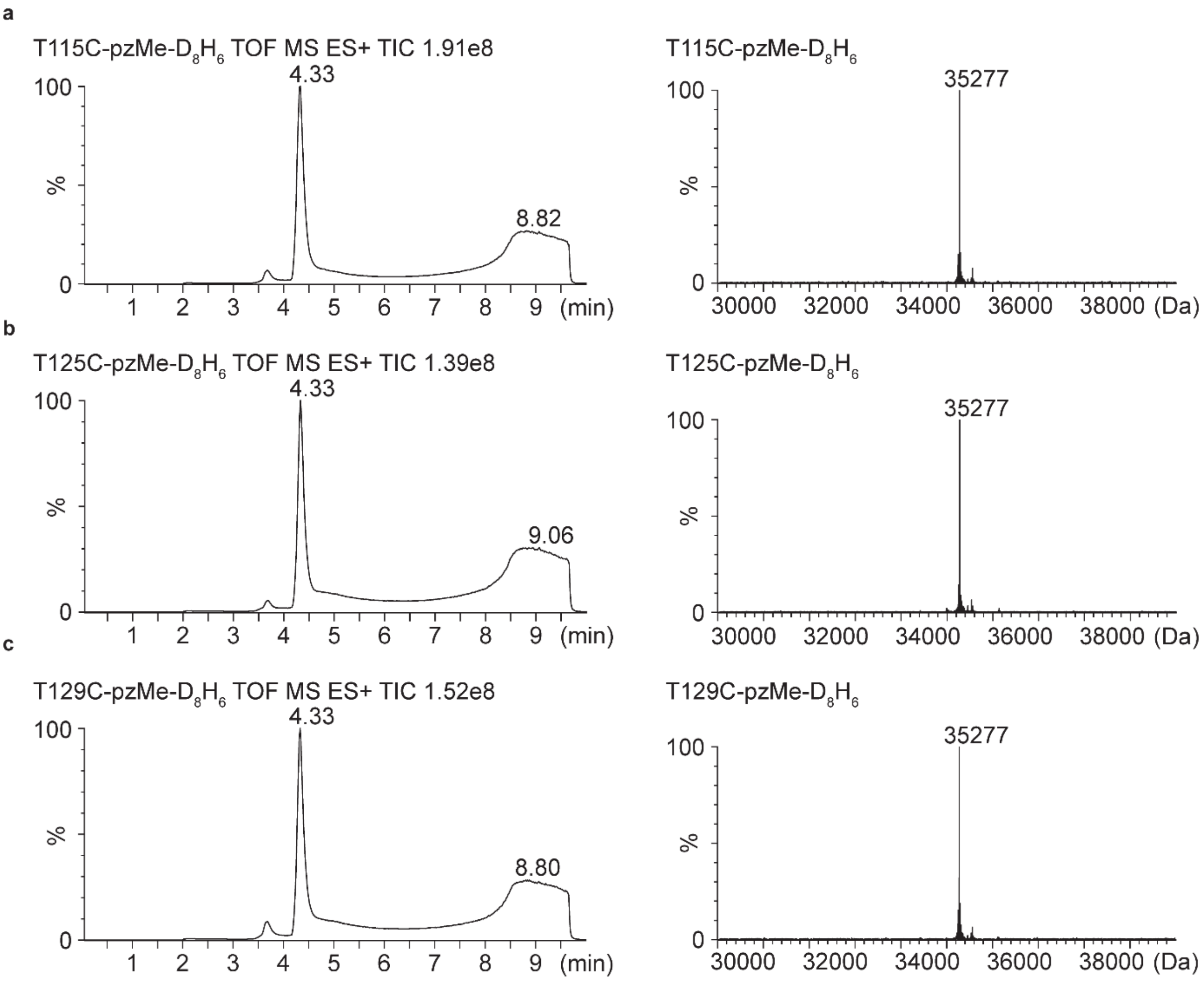

**Fig. 4 | LC–MS characterization of photopores obtained by modification with Br-pzMe.** The left panels are total ion current chromatograms. The right panels are deconvoluted ESI–MS spectra in the range of 30-40 kDa. **a,** T115C-pzMe-$D_8H_6$ monomer: mass = 35277 (calculated) and 35277 (observed). **b,** T125C-pzMe-$D_8H_6$ monomer: mass = 35277 (calculated) and 35277 (observed). **c,** T129C-pzMe-$D_8H_6$ monomer: mass = 35277 (calculated) and 35277 (observed).

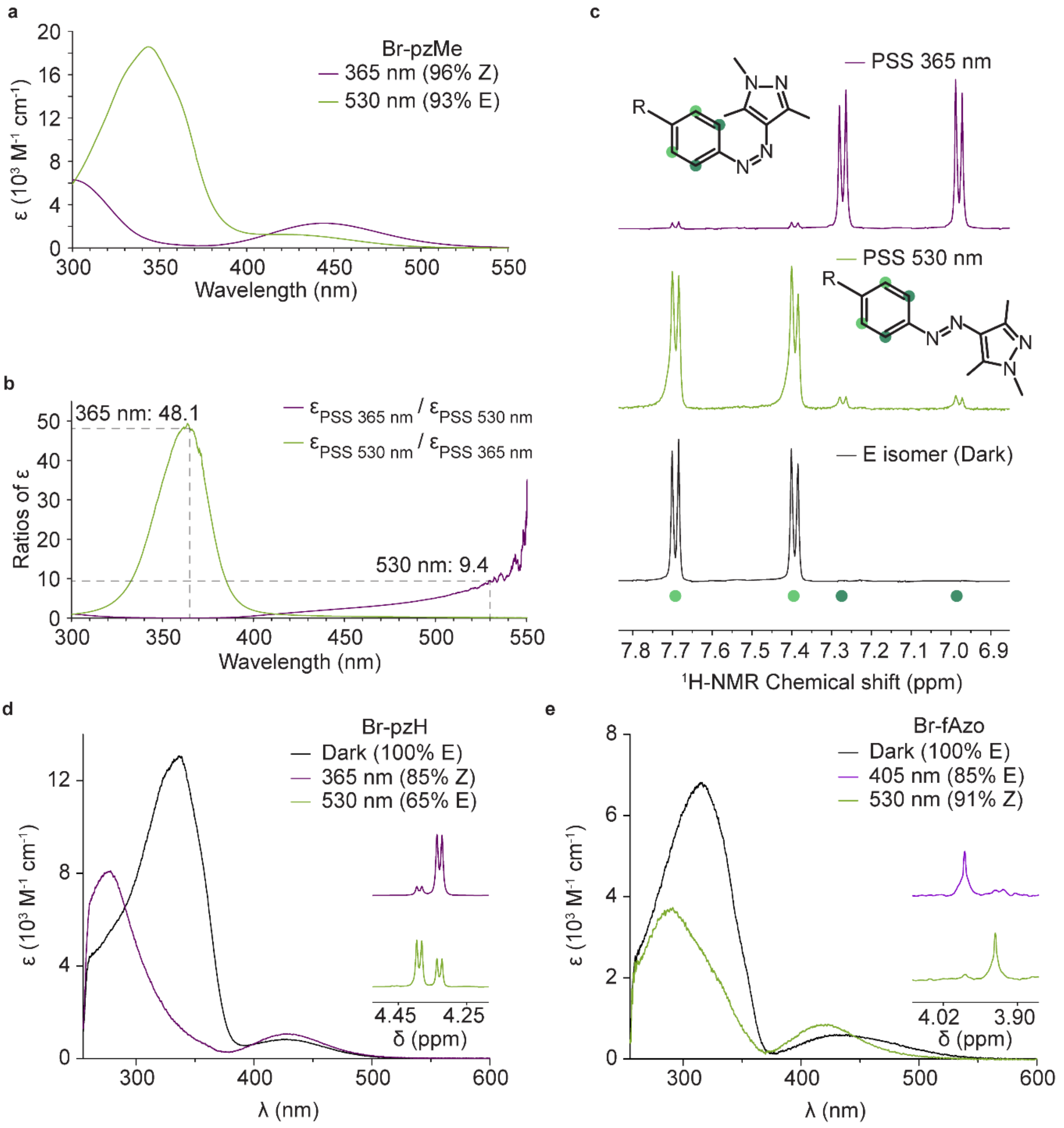

**Fig. 5 | Quantitative photoswitching of Br-pzMe. a,** UV/Vis absorption spectra of Br-pzMe in photostationary states (PSS) in DMSO. **b,** The ratio of Br-pzMe extinction coefficients (ε) between 365 and 530 nm calculated from the data in **a**. The higher ε of the E isomer of Br-pzMe at 365 nm biased the equilibrium towards the Z isomer at PSS 365 nm. Similarly, the Z isomer of pzMe exhibited higher ε at 530 nm, which favored isomerization to the E isomer at PSS 530 nm. **c,** $^{1}$H NMR signal of the aryl protons (green) of Br-pzMe in DMSO-$d_6$. **d,** UV/Vis absorption spectra of Br-pzH in the dark after heating and at PSS 365 nm. The right panel shows parts of the $^{1}$H NMR spectra of Br-pzH at PSS 365 and 530 nm. **e,** UV/Vis absorption spectra of Br-fAzo in the dark after heating and at PSS 530 nm. The right panel shows the parts of the $^{1}$H NMR spectra of Br-fAzo at PSS 405 and 530 nm. The UV/Vis spectra and $^{1}$H NMR were recorded in DMSO-$d_6$ at 298 K.

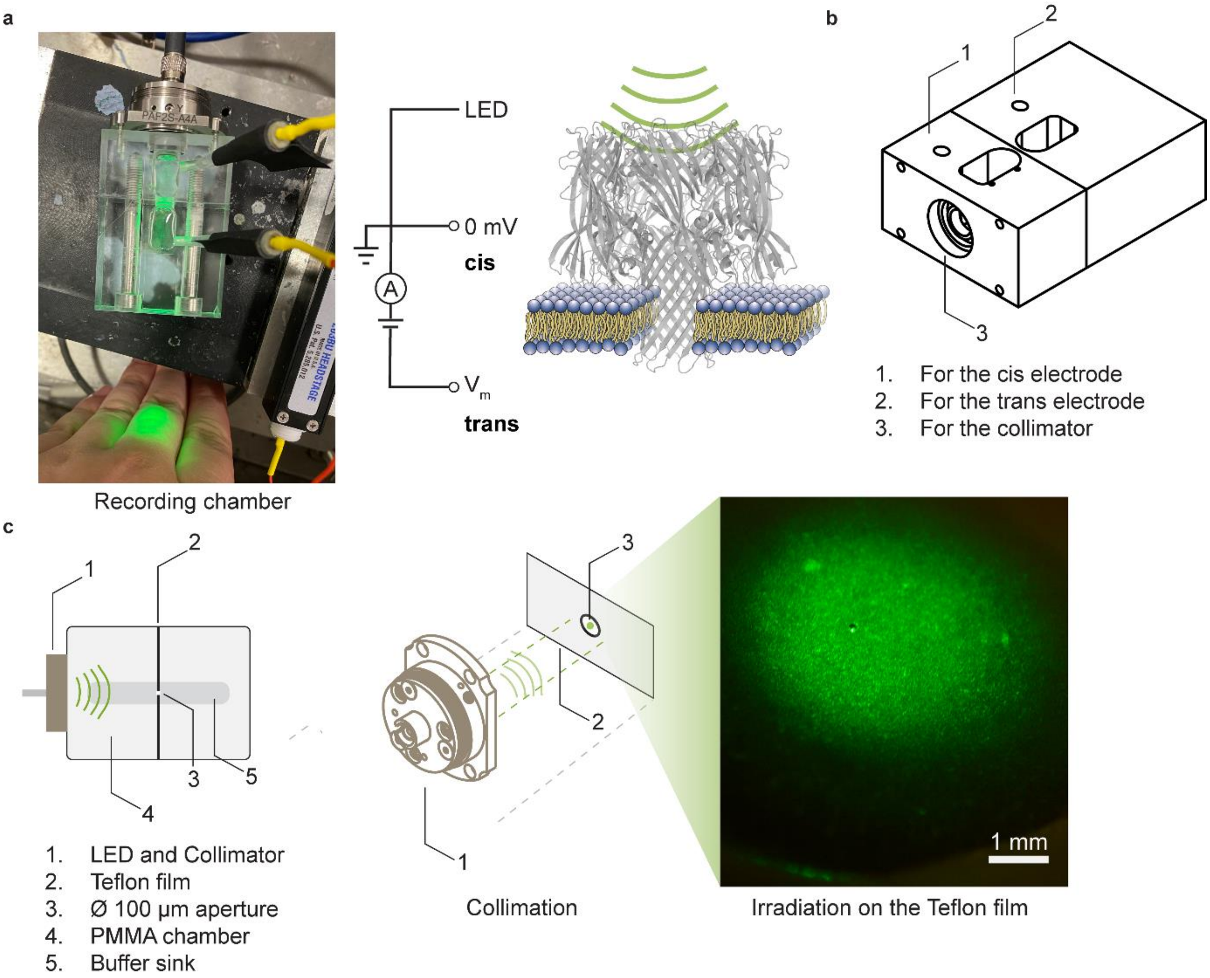

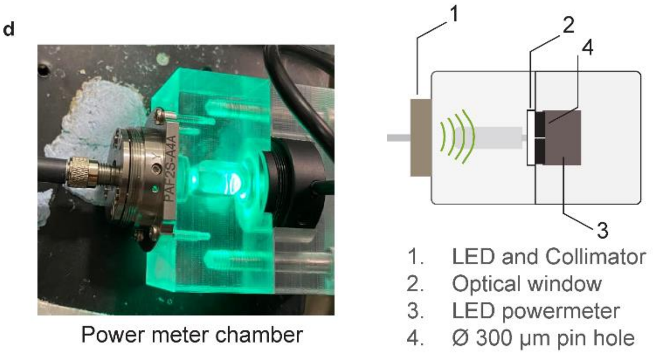

e

| Irradiance of LED on the planar lipid bilayer | | |
|---|---|---|
| Wavelength (nm) | High power (mW/cm$^2$) | Low power (mW/cm$^2$) |
| 365 | 36.2 | 13.3 |
| 405 | 75.5 | 28.7 |
| 455 | 126.2 | 56.3 |
| 530 | 30.3 | 14.6 |

**Fig. 6 | Custom recording chamber for planar lipid bilayer (PLB) experiments under irradiation. a,** A recording chamber for photopore characterization. An electrical potential was applied across the PLB with Ag/AgCl electrodes (yellow wires). The cis electrode was grounded, and the compartment with the trans electrode was connected to an optical fiber (Ø 1000 µm, NA = 0.39). **b,** Technical schematic of the chamber. The front hole was attached to a collimator and an optical window (not shown). Electrodes were inserted into buffer-containing compartments contained 1 mL buffer each. **c,** Components of the chamber. The fiber-coupled LED projected light to a Teflon film between the compartments. When adjusted, a collimated beam was projected onto the aperture (Ø ~100 µm). **d,** Measurement of light intensity. The irradiance of the light was measured with a power meter by replacing the trans part of the chamber. The collimated light was masked by a Ø 300 µm pinhole. The beam profile through the pinhole was approximately the same as that through the aperture in the Teflon. **e,** Irradiance values of LEDs measured at 365 nm, 405 nm, 455 nm and 530 nm. The irradiance was calculated from the intensity through the pinhole. Irradiance (Q) = Intensity / Pinhole area.

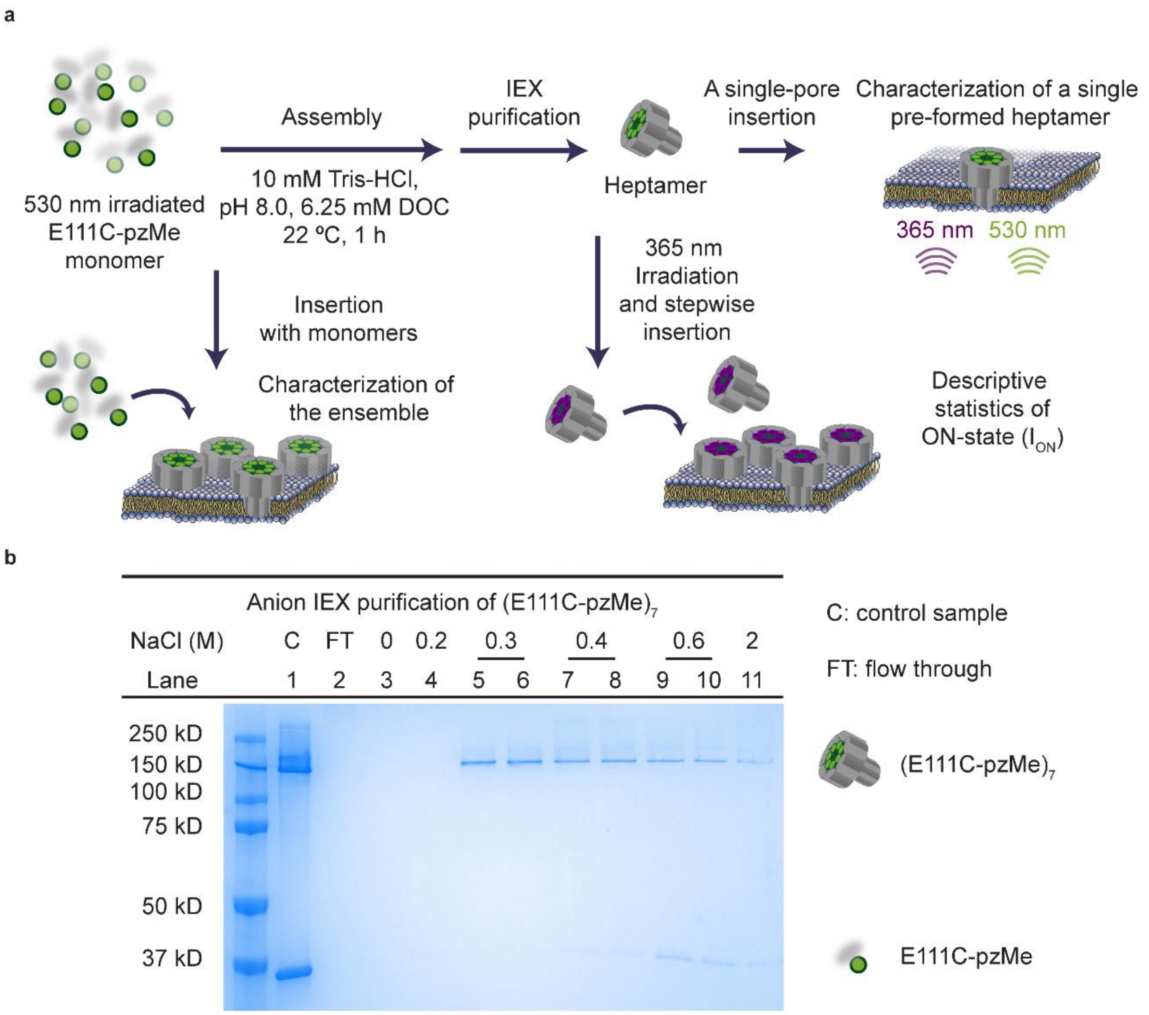

**Fig. 7 | Photopore preparation and characterization.** $(E111C\text{-}pzMe)_7$ is described as an example, but the procedures apply to other photopores. **a,** To obtain homoheptamers, monomers were incubated with sodium deoxycholate (DOC) buffer after 530 nm irradiation. Remaining monomer was removed by anion exchange chromatography (IEX). The purified homoheptamer was examined by electrical recording at the single-molecule level. A population of ON-state currents ($I_{ON}$) was examined by observing the stepwise insertion of homoheptamers generated with DOC and irradiated at 365 nm. The OFF-state current was not pursued as each pore only contributed few pA, which was indistinguishable from noise. For ensemble characterization, monomers irradiated at 530 nm were added directly to the recording chamber. **b,** Monitoring the IEX purification by gel electrophoresis. Each monomer had a $D_8H_6$ tag. The homoheptamer in lanes 5 and 6 was used for recording. Anion IEX conditions: 20 mM Tris-HCl, pH 7.5, NaCl gradient 0.2→0.6 M. Gel: 10% SDS-PAGE with Tris-Glycine SDS buffer.

a

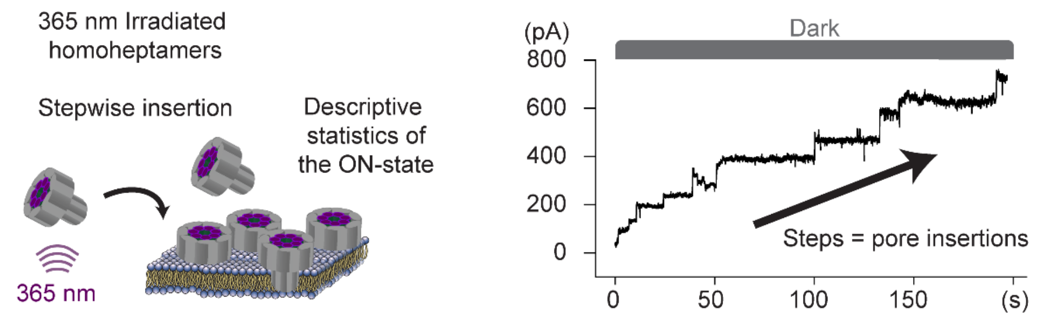

b

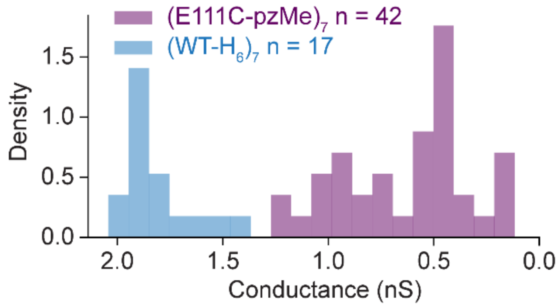

c

| Conductance of ON-state by stepwise insertion | |
|---|---|
| Species | g (nS) |
| $(E111C\text{-}pzMe)_7$ | 0.62 ± 0.30 |
| $(E111C\text{-}pzMe)_7$ max | 1.27 |
| $(E111C\text{-}pzMe)_7$ min | 0.12 |
| $(WT\text{-}H_6)_7$ | 1.81 ± 0.17 |

**Fig. 8 | Stepwise insertion of $(E111C\text{-}pzMe)_7$ irradiated at 365-nm. a,** Insertion of $(E111C\text{-}pzMe)_7$ in the $(0^E,7^Z)$ state in the dark. $(E111C\text{-}pzMe)_7$ was irradiated at 365 nm for 30 minutes and then introduced into the cis compartment. **b,** Histogram of the current steps produced by pore insertions in **a**. The current units in pA were converted to conductance units in nS consistent with the recording conditions. $(WT\text{-}H_6)_7$ exhibited a mono-disperse distribution of conductance values. In contrast, the distribution of $(E111C\text{-}pzMe)_7$ conductance values was broader. **c,** Pore conductance values. The mean unitary conductance of $(WT\text{-}H_6)_7$ was three times that of $(E111C\text{-}pzMe)_7$ after 365 nm irradiation. The current trace was recorded at +100 mV using a 5 kHz in-line Bessel filter at 25 kHz sampling frequency, and a 20 Hz digital Bessel filter was used for data analysis. Recording conditions: 2 M KCl, 10 mM Tris-HCl, 0.1 mM EDTA, pH 8.5, 24 ± 1 °C.

a

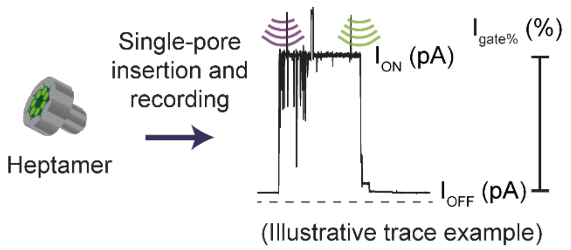

b

| ON-OFF switching of (E111C-pzMe)$_7$ Single pre-heptamerised pore insertion | | | |
|---|---|---|---|
| Repeat | $I_{ON}$ (pA) | $I_{OFF}$ (pA) | $I_{gate\%}$ (%) |
| 1 | 98.7 | 1.5 | 98.5 |
| 2 | 98.5 | 4.1 | 95.8 |
| 3 | 95.6 | 8.7 | 90.9 |
| 4 | 93.4 | 2.5 | 97.4 |
| 5 | 92.8 | 3.4 | 96.3 |
| 6 | 85.7 | 2.7 | 96.8 |
| ● 7 | 64.0 | 1.7 | 97.4 |
| 8 | 55.5 | 1.3 | 97.7 |
| 9 | 50.1 | 4.3 | 91.4 |
| 10 | 49.7 | 3.2 | 93.6 |
| 11 | 49.3 | 3.9 | 92.1 |
| 12 | 48.8 | 5.1 | 89.5 |
| 13 | 40.8 | 3.0 | 92.5 |
| 14 | 26.0 | 0.9 | 96.4 |
| 15 | 24.5 | 1.4 | 94.3 |
| 16 | 21.9 | 0.5 | 97.7 |
| 17 | 20.6 | 1.6 | 92.3 |
| 18 | 15.9 | 0.5 | 96.9 |
| 19 | 14.0 | 0.6 | 95.7 |
| 20 | 13.3 | 0.5 | 96.2 |
| 21 | 9.9 | 0.4 | 96.0 |
| $I_{gate\%}$ (%) = 95.0 ± 2.5 | | | |

c

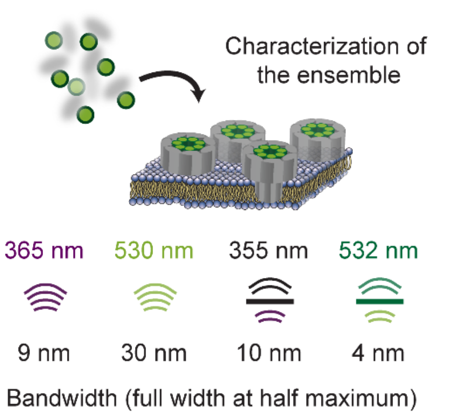

d

| ON-OFF switching of (E111C-pzMe)$_7$ ensemble | | | |
|---|---|---|---|
| Number of pores | $I_{ON}$ (pA) | $I_{OFF}$ (pA) | $I_{gate\%}$ (%) |
| 120 | 7692 | 1167 | 84.8 |
| 106 | 6808 | 961 | 85.9 |
| 113 | 7256 | 892 | 87.8 |
| 46 | 2977 | 550 | 81.5 |
| 161 | 10297 | 1649 | 84.0 |
| $I_{gate\%}$ (%) = 84.8 ± 2.1 | | | |

e

| ON-OFF switching of (E111C-pzMe)$_7$ ensemble with in-line filter | | | |
|---|---|---|---|
| Number of pores | $I_{ON}$ (pA) | $I_{OFF}$ (pA) | $I_{gate\%}$ (%) |
| 535 | 34259 | 5421 | 84.2 |
| 337 | 21585 | 3639 | 83.1 |
| 706 | 45225 | 6986 | 84.6 |
| 173 | 11057 | 2178 | 80.3 |
| $I_{gate\%}$ (%) = 83.0 ± 1.7 | | | |

**Fig. 9 | ON-OFF switching of (E111C-pzMe)$_7$. a,** Illustrative scheme for the single-channel recording of pre-heptamerized (E111C-pzMe)$_7$ pore. ON-state current ($I_{ON}$), OFF-state current ($I_{OFF}$) and percentage gated current ($I_{gate\%}$) were determined by the current traces recorded with pre-formed homoheptamer. **b,** ON-OFF switching performance of individual pre-formed homoheptamers. Each repeat refers to a new pore, and the data are sorted with descending $I_{ON}$. Despite the diverse values of $I_{ON}$ (0.51 ± 0.31 nS, n = 21 repeats), which are consistent with the stepwise insertion results, the mean value of $I_{gate\%}$ is more tightly defined. The seventh repeat was with the pore used to collect the 40-minute recording of ON-OFF switching (**Fig. 2b**). **c,** Illustrative scheme for an (E111C-pzMe)$_7$ ensemble generated by the insertion of monomers. The ensemble was characterized with a pair of in-line filters to shift the LED irradiation wavelengths from 365/530 nm to 355/532 nm. The filter also narrowed the spectral bandwidth from 9/30 nm to 10/4 nm while reducing the output intensity. **d-e,** ON-OFF switching of the ensemble without and with the filters. The current and $I_{gate\%}$ values were calculated from traces recorded at +100 mV using a 5 kHz in-line Bessel filter at 25 kHz sampling frequency. Recording conditions: 2 M KCl, 10 mM Tris-HCl, 0.1 mM EDTA, pH 8.5, 24 ± 1 °C.

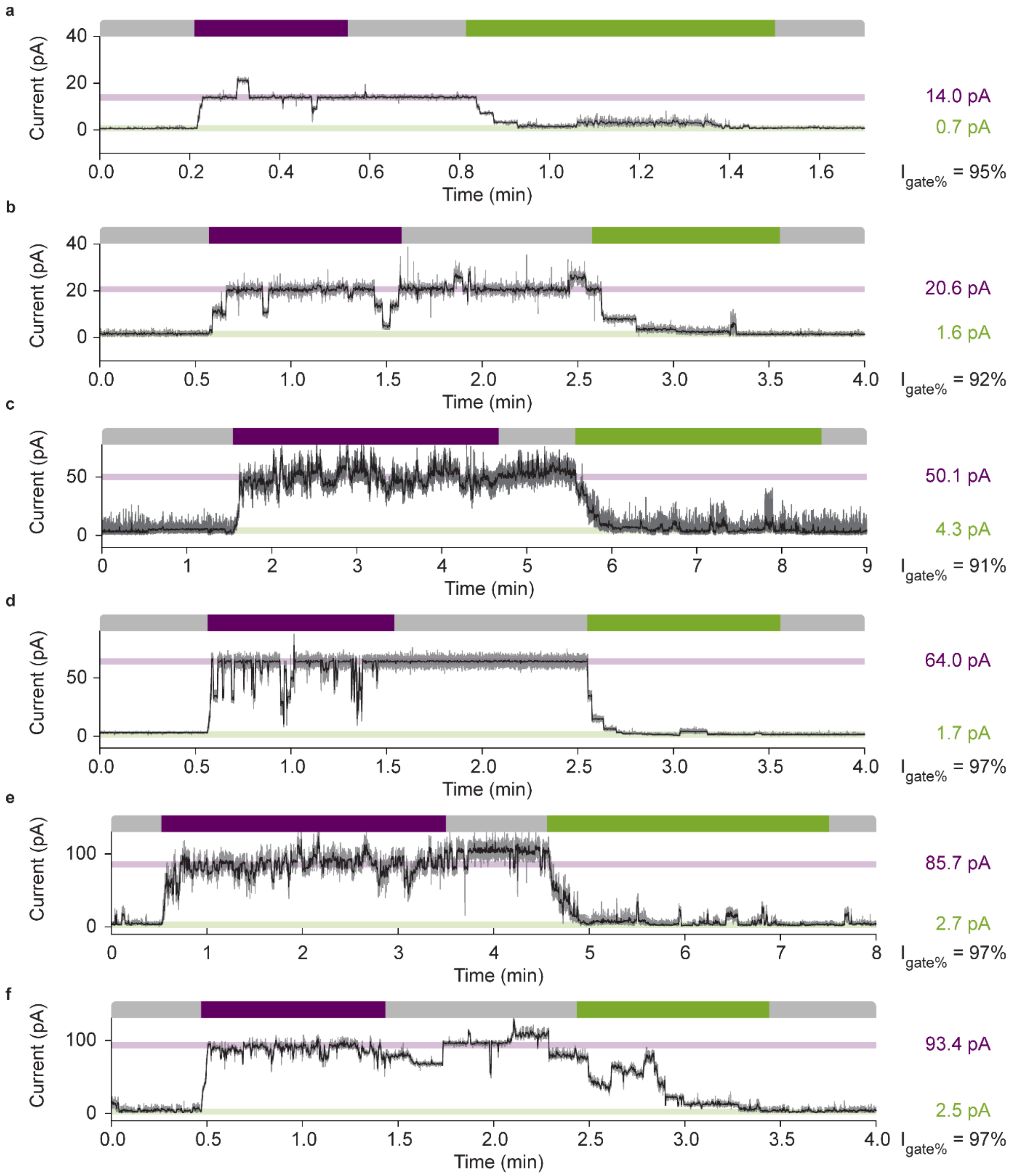

**Fig. 10 | Representative determinations of $I_{gate\%}$ values for single (E111C-pzMe)$_7$ pores.** The moving averages of the ionic currents (grey traces) were generated with a Savitzky–Golay filter and are shown in black. The top bar shows the irradiation cycle of dark→365 nm→dark→530 nm→dark. The $I_{gate\%}$ of all traces is >90%, suggesting that $I_{gate\%}$ appears to be consistent regardless of the absolute current ($I_{ON}$ and $I_{OFF}$). The current traces were recorded at +100 mV using a 5 kHz in-line Bessel filter at 25 kHz sampling frequency, and a 20 Hz digital Bessel filter was used for data analysis. Recording conditions: 2 M KCl, 10 mM Tris-HCl, 0.1 mM EDTA, pH 8.5, 24 ± 1 °C.

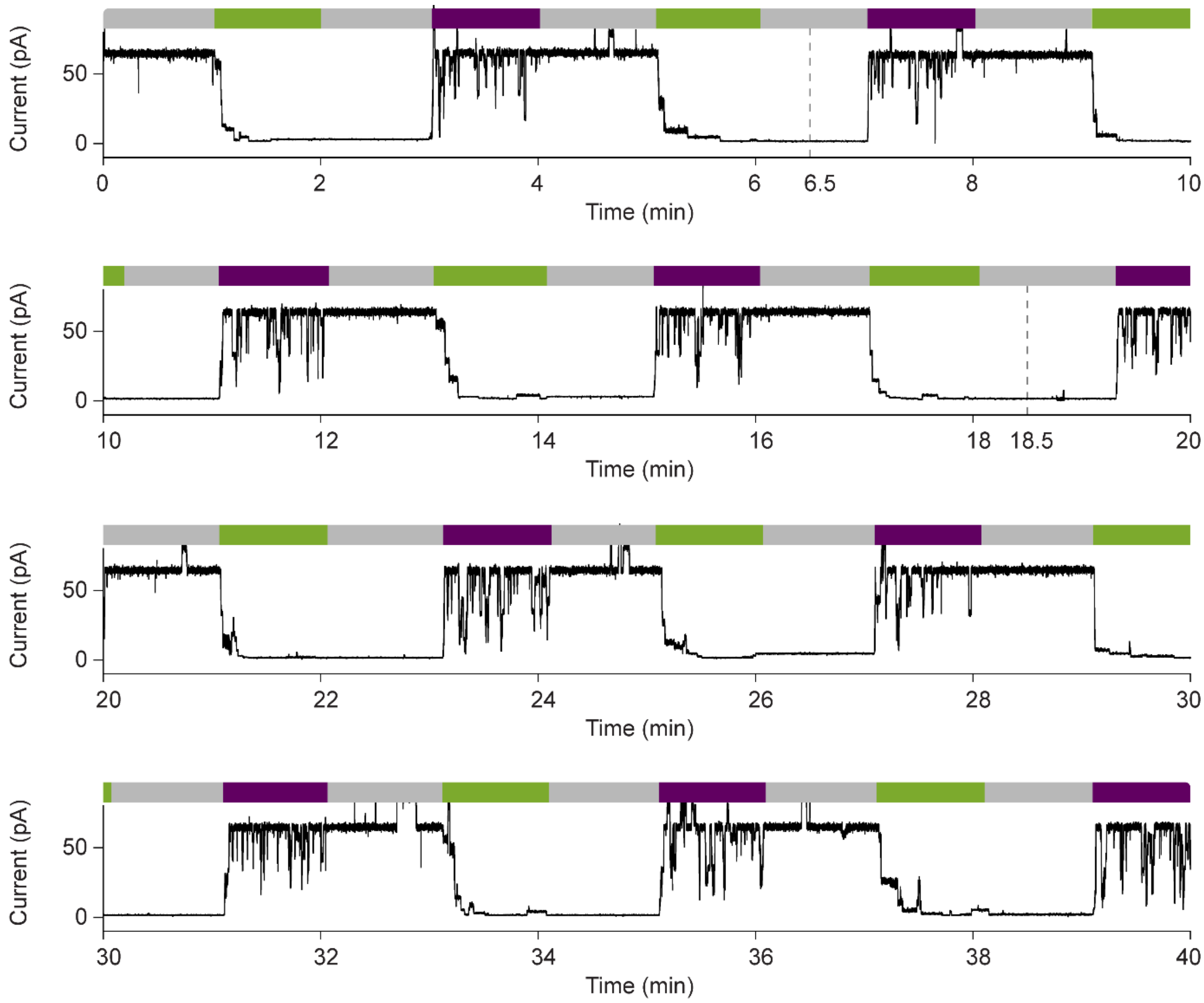

**Fig. 11 | Prolonged trace of ON-OFF cycles recorded with a single (E111C-pzMe)$_7$ pore.** Ten cycles were recorded over 40 minutes. **Fig. 2d** was taken from the region between the two grey dashed lines at 6.5 and 18.5 min. The current trace was recorded at +100 mV using a 5 kHz in-line Bessel filter at 25 kHz sampling frequency, and a 20 Hz digital Bessel filter was used for data analysis. Recording conditions: 2 M KCl, 10 mM Tris-HCl, 0.1 mM EDTA, pH 8.5, 24 ± 1 °C.

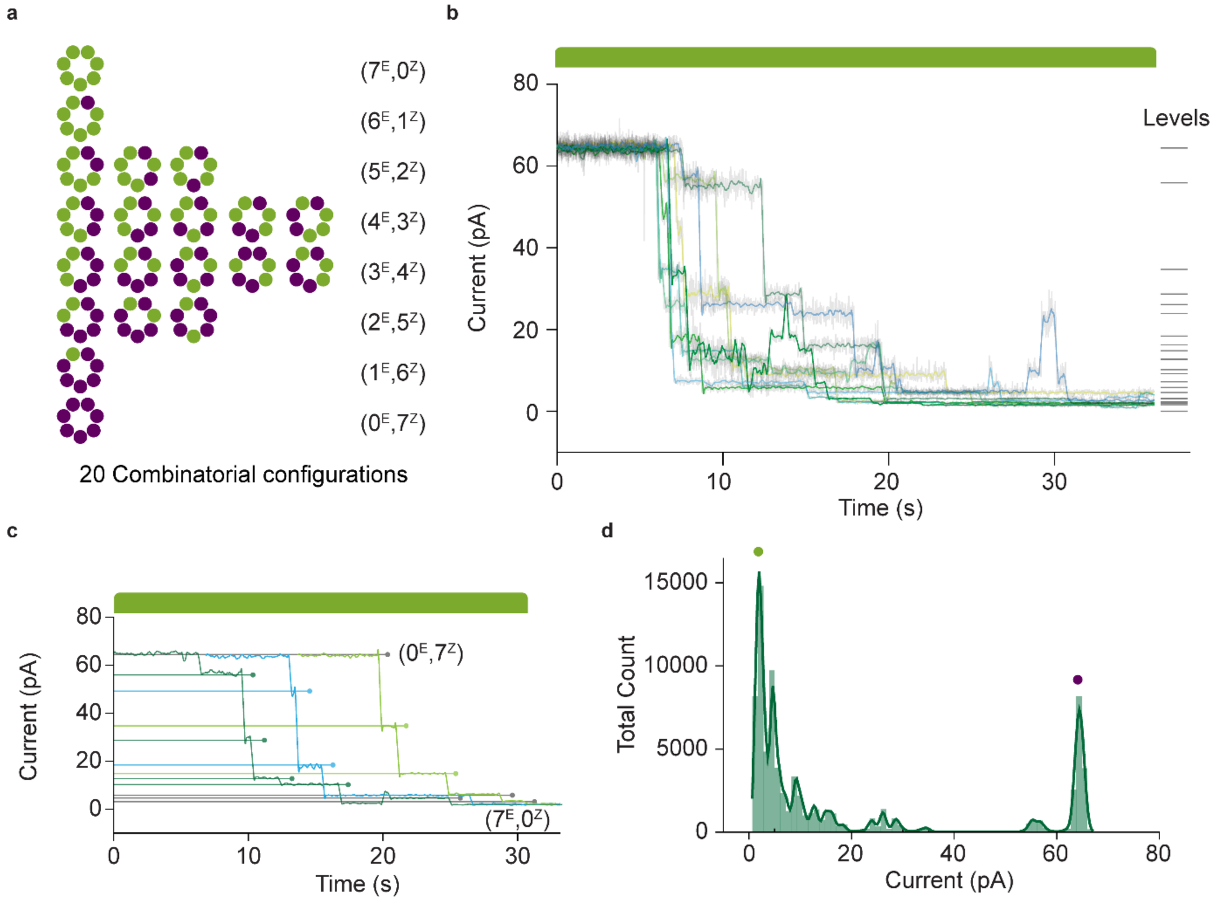

**Fig. 12 | Stochastic behavior of pzMe photoisomerization within a single (E111C-pzMe)$_7$ pore. a,** Representation of 20 combinatorial configurations of pzMe in a pore. The origins of the heterogeneity in unitary conductance of the (E111C-pzMe)$_7$ pore lies in the relative population and positioning of the E/Z pzMe molecules, and the stacking of the pzMe molecules in the axial-chiral β barrel. **b,** Overlay of ten Z-to-E transitions extracted from a current trace of a single (E111C-pzMe)7 pore. All transitions consistently and reproducibly started from the 64 pA level (ON) and ended at 1.6 pA (OFF), but the intermediate steps were stochastic. **c,** Three traces selected from **b** to illustrate the non-overlapping intermediate current levels. In particular, multiple levels were derived from the ($0^E$,$7^Z$) level, suggesting that several current levels can be ascribed to the ($1^E$,$6^Z$) configuration. **d,** Histogram of the current levels presented in **b**. The green and purple dots are the OFF and ON states.

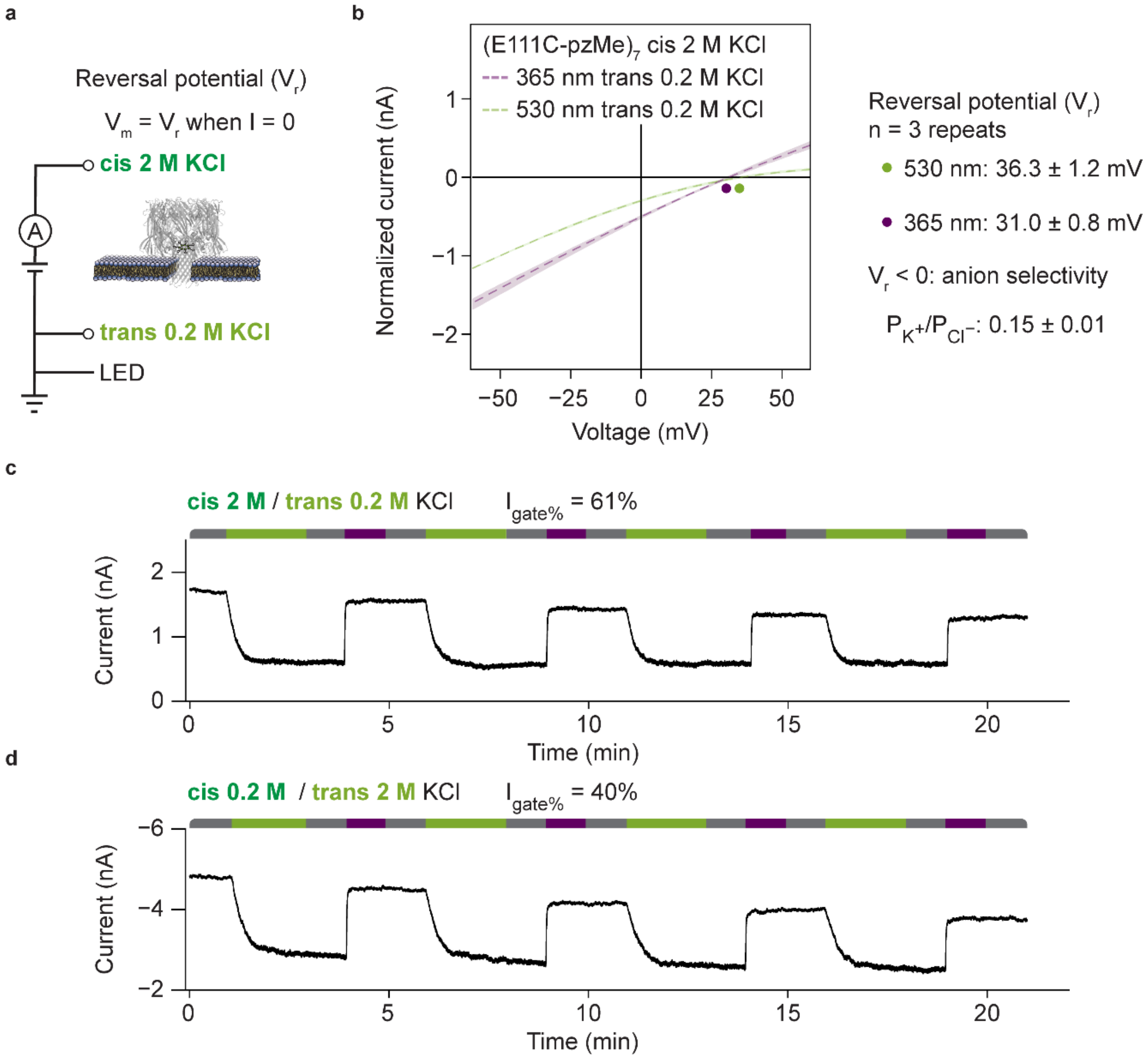

**Fig. 13 | Charge selectivity of (E111C-pzMe)$_7$. a,** Transmembrane potential generated by an asymmetric KCl concentration across the lipid bilayer. The reversal potential ($V_r$) is defined as the potential required to bring the current flow to zero. **b**, I–V characterization of an ensemble of (E111C-pzMe)$_7$. The reversal potential was measured after 530- and 365-nm irradiation. The shading around the curves represents the 95% confidence interval after three repeats. The measured $V_r$ suggests anion selectivity. **c-d,** Currents driven by a KCl concentration gradient. The values of $I_{gate\%}$ in **c** and **d** was consistent with the voltage-dependence of $I_{gate\%}$ reported in **Fig. 3c**. The experiment was recorded at 25 kHz sampling frequency and filtered using a 5 kHz in-line Bessel filter. Recording conditions: (**b**, **c**) 2 M / 0.2 M KCl, (**d**) 0.2 M / 2 M KCl, 10 mM Tris-HCl, 0.1 mM EDTA, pH 8.5, 24 ± 1 °C.

### Synthesis and characterization of photoswitches

#### Materials and methods

All reagents and solvents were purchased from commercial sources and used without further purification. Where necessary, solvents were dried by passing through an MBraun MPSP-800 column and degassed with nitrogen. Triethylamine was distilled from and stored over potassium hydroxide. Column chromatography was carried out on Merck® silica gel 60 under a positive pressure of nitrogen. Where mixtures of solvents were used, ratios were reported by volume. NMR spectra were recorded on a Bruker AVIII 400, Bruker AVII 500 (with cryoprobe), Bruker NEO 600 with broadband helium cryoprobe and Bruker AVIII 500 spectrometers. Chemical shifts were reported as δ values in ppm. Mass spectra were carried out on a Waters Micromass LCT and Bruker microTOF spectrometers. UV/Vis spectra were recorded on a V-770 UV/Visible/NIR Spectrophotometer equipped with Peltier temperature controller and stirrer using quartz cuvettes of 1 cm path length. Experiments were conducted at 25°C unless otherwise stated.

#### PSS Determination and UV/Vis spectra

UV/Vis spectra were determined in the DMSO-$d_6$ solution. Extinction coefficients were determined by recording UV/Vis spectra for the *E* isomer at 10, 20, 30, 40 µM in DMSO. The absorbance at the maximum of the π - π* transition of the *E*-isomers was plotted against concentration (Beer-Lambert plot) to determine the molar extinction coefficient ε. For each compound, the *E* isomer sample at 40 µM was irradiated with the desired wavelength of light to generate a photo-stationary state and another spectrum was run. This spectrum was normalized to units of ε and overlaid with the dark (100% *E* isomer) spectrum. Photo-irradiation of liquid samples was carried out using Thorlabs high-power mounted LEDs (models M530L4 (green, 530 nm), M365L2 (UV, 365 nm)) and in-house custom built set-ups using optical components supplied by Thorlabs. Photo-stationary states (PSS) were determined using $^1$H NMR spectroscopy.

#### Abbreviations

AcOH: Acetic Acid; Boc: tert-butyloxycarbonyl; DCM: Dichloromethane; DIPA: N,N-Diisopropylamine; DIPEA: N,N-Diisopropylethylamine; DMAP: 4-dimethylaminopyridine; HRMS: High resolution mass spectrometry; MeCN: Acetonitrile; MeOH: Methanol; PhMe: Toluene; Phth: Phthaloyl; PSS: Photo-stationary state; rt: Room temperature; TFA: Trifluoroacetic acid; THF: Tetrahydrofuran; TMS: trimethylsilyl

#### Synthesis routes

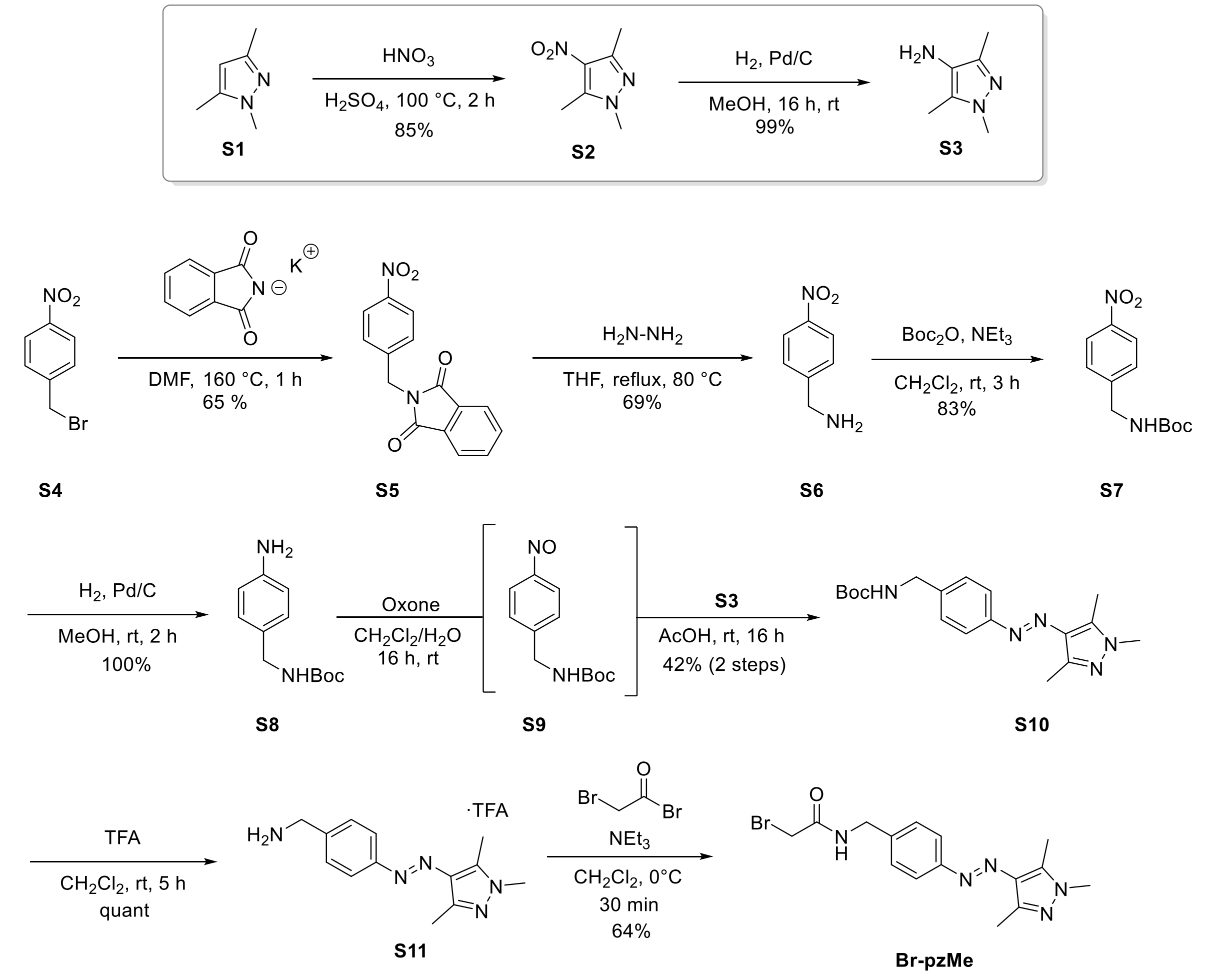

**Scheme 1 |** Preparation of **Br-pzMe**. The synthesis commenced from commercially available **S1**, which was nitrated to **S2** and subsequently hydrogenated to **S3**. The azobenzene scaffold was generated by treatment of **S4** with potassium phthalimide to afford **S5**, which was deprotected to **S6**, reprotected to **S7** and finally hydrogenated to **S8**. This was subjected to a Mills coupling reaction *via* **S9**, which was reacted with **S3** to generate azobenzene **S10** in moderate yields. Boc deprotection to **S11** followed by treatment with bromoacetyl bromide gave rise to **Br-pzMe** in good yields.

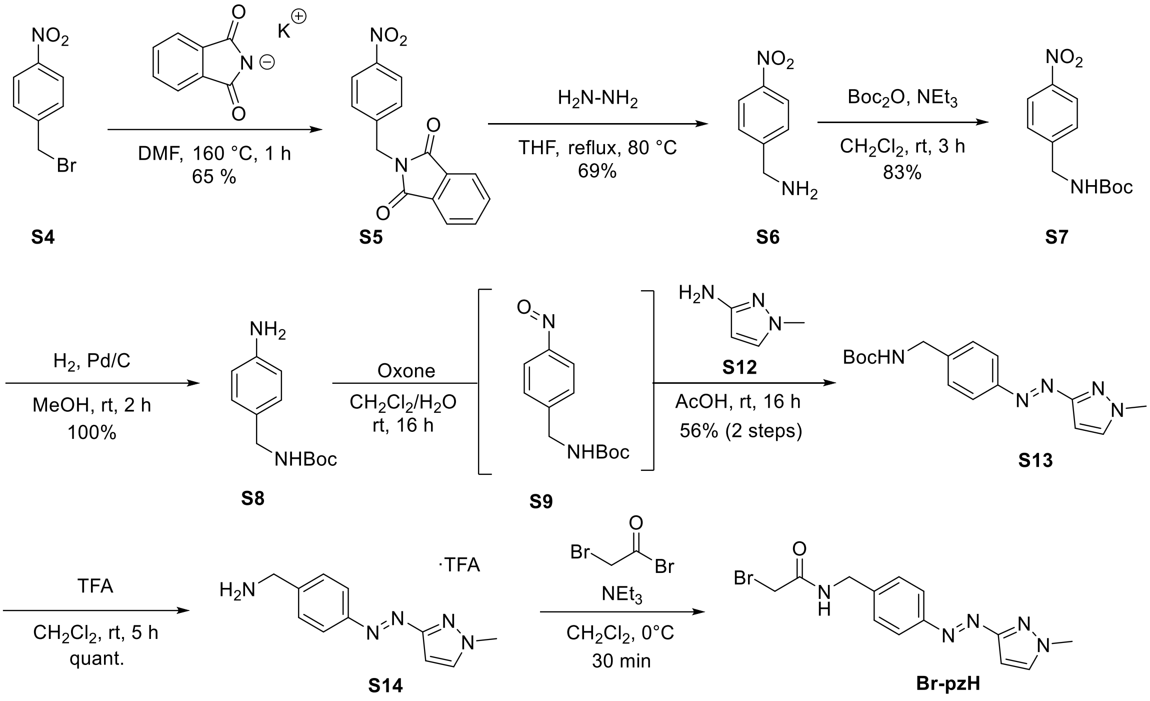

**Scheme 2 |** The synthesis commenced from commercially available **S4**, which was nitrated to **S5** and subjected to hydrazinolysis to **S6**, reprotected to **S7** and finally hydrogenated to **S8**. This was subjected to a Mills coupling reaction *via* **S9**, which was reacted with **S12** to generate azobenzene **S14** in good yields. Boc deprotection to **S14** followed by treatment with bromoacetyl bromide gave rise to **Br-pzH** in good yields.

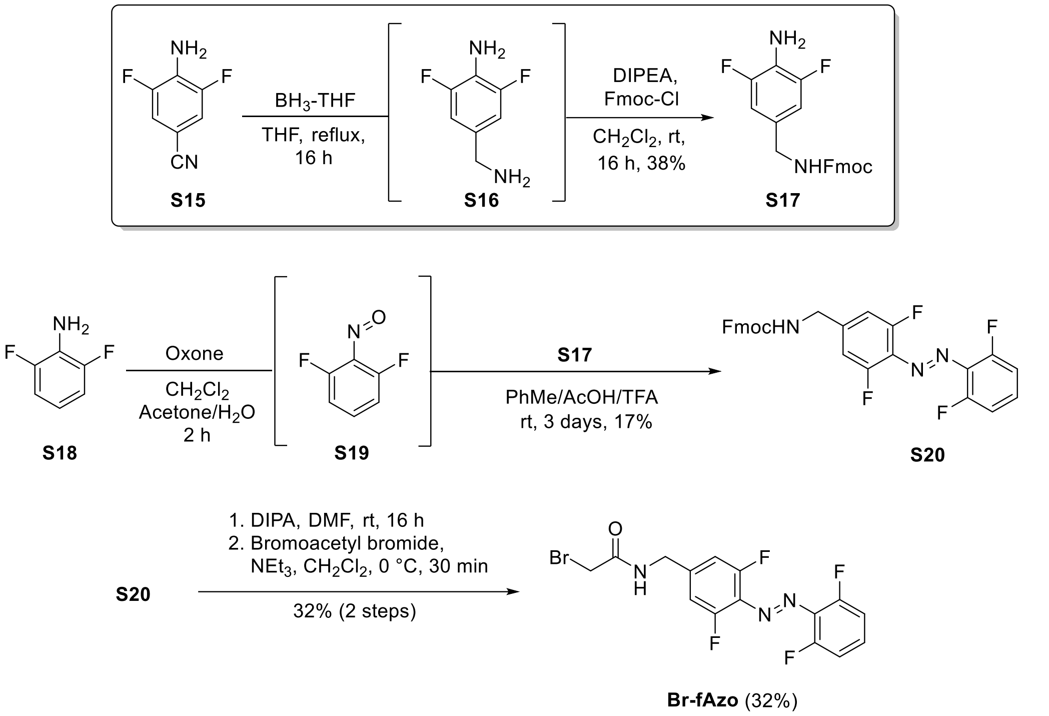

**Scheme 3 |** Preparation of **fAzo** derivatives. **S15** was reduced to unstable **S16**, followed by Fmoc-protection to **S17**. This was azo coupled to **S18** to generate **S20**, which was deprotected using DIPA and treated with bromoacetyl bromide to afford the **Br-fAzo** derivatives.

#### Synthesis and characterization

#### Synthesis of Br-pzMe

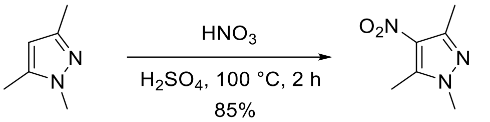

S1 S2

Nitro compound **S2**. Prepared according to a modified literature procedure.[5] Pyrazole **S1** (1.5 g, 13.6 mmol) was dissolved in sulfuric acid (7.5 mL). The solution was cooled to 0 °C, and then nitric acid (6 mL, 150 mmol, 11 eq) was added. The solution was heated to 100 °C for 2 hours. The mixture was cooled to room temperature, then poured into ice water. The mixture was slowly basified under ice cooling with NaOH pellets. The precipitated solid was collected via vacuum filtration, washed with water, and dried to afford the title compound as an off-white solid (1.8 g, 11.6 mmol, 85%). $^1$H NMR (400 MHz, $CDCl_3$) δ 3.77 (s, 3H), 2.61 (s, 3H), 2.50 (s, 3H). Data were consistent with that given in the literature[5].

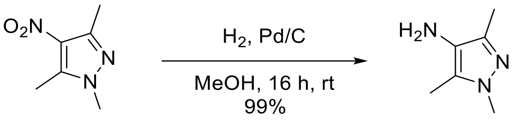

S2 S3

Aniline **S3**. Prepared according to a modified literature procedure.[6] Nitro compound **S2** (1.8 g, 11.6 mmol) was dissolved in MeOH (25 mL) and placed under nitrogen. Palladium 10% on carbon (250 mg) was added, then the reaction was stirred under hydrogen atmosphere for 16 hours at room temperature. The reaction was filtered over celite and washed with $CH_2Cl_2$. The filtrate was concentrated to afford the title compound as a yellow solid. (1.44 g, 11.5 mmol, 99%). $^1$H NMR (400 MHz, $CDCl_3$) δ 3.66 (s, 3H), 2.46 – 2.31 (br s, 2H), 2.15 (s, 3H), 2.13 (s, 3H). Data were consistent with that given in the literature.[6]

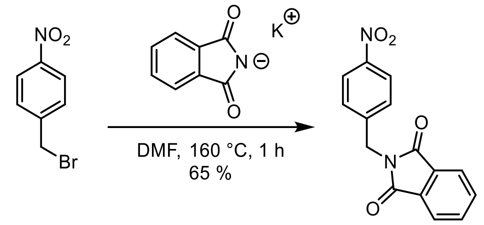

S4 S5

Phthalimide **S5**. Prepared according to a modified literature procedure.[7] *p*-nitrobenzyl bromide **S4** (7 g, 32.40 mmol) was dissolved in DMF (45 mL). Potassium phthalimide (6.3 g, 34 mmol, 1.05 eq) was added and the reaction was stirred at reflux temperature for 1 h. The reaction was cooled, and ice water was added. The filtrate was dissolved in hot EtOAc (300 mL) and allowed to recrystallize for two days at room temperature in a 500 mL flask. The solid was collected and washed with cold EtOAc to afford the title compound as white crystals (5.9 g, 20.9 mmol, 65%). $^1$H NMR (400 MHz, $CDCl_3$) δ 8.22 – 8.15 (m, 2H), 7.88 (dd, $J$ = 5.4, 3.1 Hz, 2H), 7.75 (dd, $J$ = 5.5, 3.1 Hz, 2H), 7.62 – 7.56 (m, 2H), 4.93 (s, 2H). Data consistent with that given in the literature.[7]

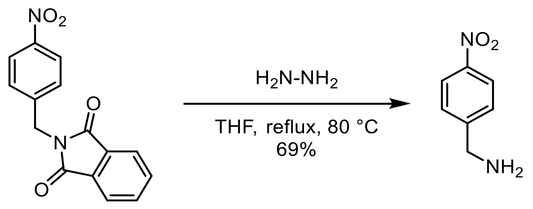

**S5** **S6**

Amine **S6**. Prepared according to a modified literature procedure.[8] Nitro compound **S5** (10 g, 35.43 mmol) was dissolved in THF (100 mL). Hydrazine hydrate (8 mL) was added and the reaction was stirred at reflux temperature for 16 hours. The reaction was concentrated, then taken up in EtOAc. The suspension was filtered over cotton, and the filtrate washed with 1 M NaOH. The organic layer was dried, then concentrated to afford the title compound as a yellow oil (3.7 g, 24.3 mmol, 69%). $^1$H NMR (400 MHz, $CDCl_3$) δ 8.23 – 8.16 (m, 2H), 7.50 (d, $J$ = 8.3 Hz, 2H), 4.01 (s, 2H). Data were consistent with that given in the literature.[8]

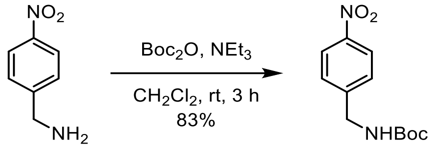

**S6** **S7**

Nitro compound **S7**. Prepared according to a modified literature procedure.[9] Amine **S6** (3.7 g, 24.32 mmol) was dissolved in $CH_2Cl_2$ (50 mL). $NEt_3$ (10.2 mL, 73 mmol, 3 eq) was added. Then $Boc_2O$ (8 g, 36.5 mmol, 1.5 eq) in $CH_2Cl_2$ (50 mL) was added dropwise. The reaction was stirred for 3 hours, after which it was poured onto ice water (50 mL). The reaction was recrystallized from isopropyl ether (40 mL) to afford the title compound as a yellow crystalline solid (5.1g, 20.2 mmol, 83%). $^1$H NMR (400 MHz, $CDCl_3$) δ 8.23 – 8.14 (m, 2H), 7.45 (d, $J$ = 8.6 Hz, 2H), 4.99 (s, 1H), 4.42 (d, $J$ = 6.3 Hz, 2H), 1.47 (s, 9H). Data were consistent with that given in the literature.[9]

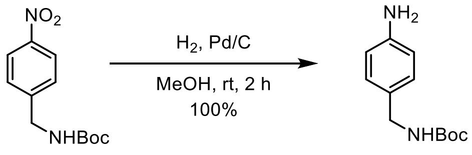

**S7** **S8**

Phthalimide **S8.** Prepared according to a modified literature procedure.[9] Nitro compound **S7** (2.42 g, 9.59 mmol) was dissolved in EtOH (40 mL). The reaction was placed under nitrogen, then 10% Pd/C (200 mg) was added. The reaction was stirred under $H_2$ atmosphere for 2 hours, filtered over celite, then concentrated to afford the title compound as a white solid (2.13 g, 9.59 mmol, 100%). $^1$H NMR (400 MHz, $CDCl_3$) δ 7.07 (d, $J$ = 8.0 Hz, 2H), 6.69 – 6.60 (m, 2H), 4.73 (s, 1H), 4.18 (d, $J$ = 5.7 Hz, 2H), 1.45 (s, 9H).[9]

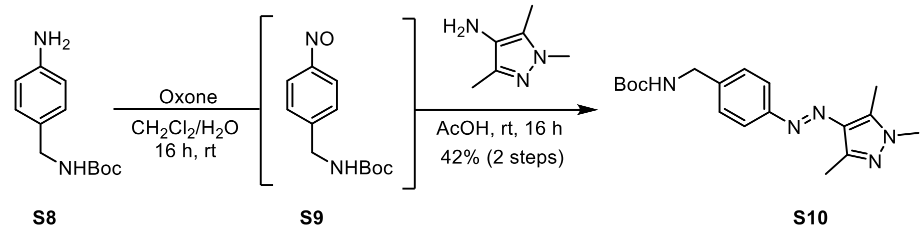

**S8** **S9** **S10**

Azobenzene **S10.** A solution of aniline **S8** (554 mg, 2.45 mmol, 1.2 equiv.) in $CH_2Cl_2$ (40 mL), was treated with a solution of Oxone® (4.9 g, 16 mmol, 6.5 equiv.) in $H_2O$ (40 mL). The biphasic reaction mixture was stirred vigorously at rt for 16 hours. The phases were separated and the aqueous phase was extracted with $CH_2Cl_2$ (20 mL). The combined organic extracts were washed with saturated $NaHCO_{3(aq)}$ solution and $H_2O$. The washed organic layer was treated with **S3** (260 mg, 2.1 mmol, 1.0 equiv.) and AcOH (30 mL). The $CH_2Cl_2$ was then removed under reduced pressure at 35 °C and the solution was stirred for 16 hours at rt. The solution was concentrated and purified by flash silica gel chromatography (0 to 10% EtOAc in $CH_2Cl_2$) to yield the title compound as a yellow solid (300 mg, 0.87 mmol, 42%). $^1$H NMR (600 MHz, $CDCl_3$) δ 7.74 (d, *J* = 8.0 Hz, 2H), 7.36 (d, *J* = 8.0 Hz, 2H), 4.88 (s, 1H), 4.36 (d, *J* = 6.0 Hz, 2H), 3.78 (s, 3H), 2.57 (s, 3H), 2.49 (s, 3H), 1.47 (s, 9H). $^{13}$C NMR (151 MHz, $CDCl_3$) δ 156.03, 153.08, 142.55, 140.31, 138.89, 135.25, 128.13, 122.15, 79.76, 44.58, 36.11, 28.56, 13.91, 10.11. HRMS-EI (m/z) Calculated for $C_{18}H_{26}N_5O_2$ $[M+H]^+$ , 344.2081; found 344.2084.

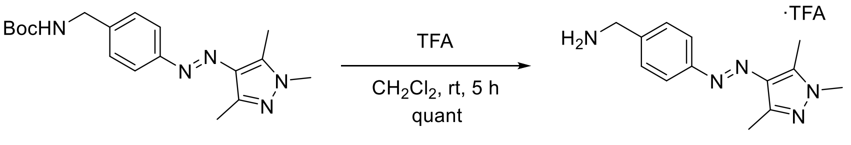

**S10** **S11**

Amine **S11.** To a solution of **S10** (100 mg, 0.29 mmol) in $CH_2Cl_2$ (5 mL) was added TFA (0.6 mL). The reaction was stirred at room temperature for 5 hours. The TFA was removed under a stream of nitrogen, then the residue was dried in vacuo to afford the title compound as a yellow solid (104 mg, 0.29 mmol, 100%). $^1$H NMR (600 MHz, MeOD) δ 7.85 – 7.81 (m, 2H), 7.58 – 7.55 (m, 2H), 4.18 (s, 2H), 3.79 (s, 3H), 2.61 (s, 3H), 2.45 (s, 3H).$^{13}$C NMR (151 MHz, MeOD) δ 155.30, 143.22, 141.59, 136.05, 135.46, 130.82, 123.36, 44.01, 36.13, 13.87, 9.76. HRMS-EI (m/z) Calculated for $C_{13}H_{18}N_5$ $[M+H]^+$ , 244.1557; found 244.1557.

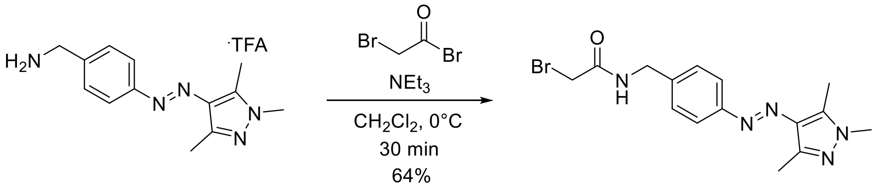

**S11** **Br-pzMe**

**Br-pzMe.** TFA salt **S11** (98 mg, 0.3 mmol) was suspended in $CH_2Cl_2$ (2.5 mL). $NEt_3$ (95 µL, 0.69 mmol, 2.5 eq) was added and the solution cooled to 0°C and bromo acetylbromide (37 µL, 0.37 mmol, 1.5 eq) was added and the solution was stirred at 0 °C for 30 minutes. The solution was diluted with $CH_2Cl_2$, then the volatiles removed by rotary evaporation (35 °C water bath). The residue was purified by flash-column chromatography (15% EtOAc in $CH_2Cl_2$) to afford the product as a yellow solid (64 mg, 175 µmol, 64%). $^1$H NMR (600 MHz, DMSO) δ 8.84 (t, *J* = 6.0 Hz, 1H), 7.69 (d, *J* = 8.0 Hz, 2H), 7.39 (d, *J* = 8.0 Hz, 2H), 4.36 (d, *J* = 6.0 Hz, 2H), 3.93 (s, 2H), 3.74 (s, 3H), 2.55 (s, 3H), 2.37 (s, 3H). $^{13}$C NMR (151 MHz, DMSO) δ 166.64, 152.55, 140.77, 140.77, 139.94, 134.81, 128.49, 121.87, 42.73, 36.42, 29.91, 14.22, 9.94. HRMS-EI (m/z) Calculated for $C_{15}H_{19}N_5OBr$ $[M+H]^+$, 364.0767; found 364.0763. HMBC showed two peaks at 140.77.

**Synthesis of Br-pzH**

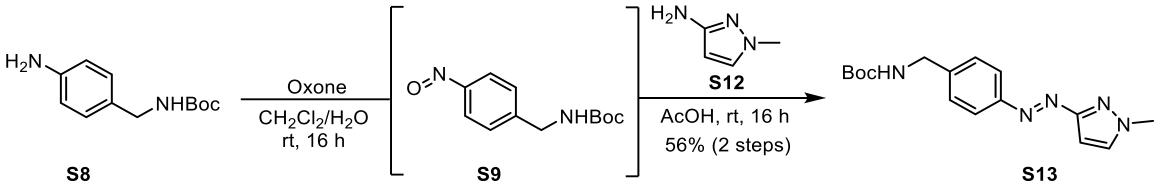

Azobenzene **S13.** A solution of aniline **S8** (452 mg, 2 mmol, 1.2 equiv.) in $CH_2Cl_2$ (37 mL), was treated with a solution of Oxone® (4 g, 13 mmol, 6.5 equiv.) in $H_2O$ (37 mL). The resulting biphasic reaction mixture was stirred vigorously at room temperature for 16 hours. Subsequently, the phases were separated and the aqueous phase was extracted with $CH_2Cl_2$ (20 mL). The combined organic extracts were washed sequentially with saturated aqueous sodium bicarbonate solution (50 mL), and $H_2O$ (50 mL). The washed organic layer containing **S9** was treated with **S12** (147 µL, 1.69 mmol, 1.0 equiv.) and AcOH (30 mL). The $CH_2Cl_2$ was then removed under reduced pressure at 35 °C and the solution was stirred for 16 hours at room temperature. The AcOH was then removed under reduced pressure. The residue was purified by flash silica gel chromatography (0 to 5% EtOAc in $CH_2Cl_2$) to yield the title compound as a yellow/brown solid (204 mg, 0.95 mmol, 56%). (*E*)-**S13**: $^1$H NMR (400 MHz, $CDCl_3$) δ 7.91 (d, *J* = 8.3 Hz, 2H), 7.45 – 7.35 (m, 3H), 6.65 (d, *J* = 2.4 Hz, 1H), 4.93 (s, 1H), 4.38 (s, 2H), 4.02 (s, 3H), 1.47 (s, 9H). $^{13}$C NMR (151 MHz, $CDCl_3$) δ 163.81, 156.06, 152.18, 142.24, 131.98, 128.13, 123.35, 96.38, 79.85, 44.54, 39.80, 28.54. HRMS-EI (m/z) Calculated for $C_{16}H_{22}N_5O_2$ $[M+H]^+$, 316.1768; found 316.1768.

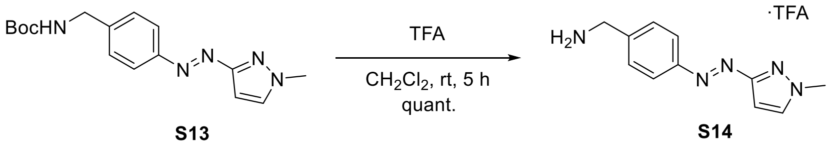

Amine **S14.** To a solution of **S13** (100 mg, 0.32 mmol) in $CH_2Cl_2$ (5 mL) was added TFA (0.6 mL). The reaction was stirred at room temperature for 5 hours. The TFA was removed under a stream of nitrogen, then the residue was dried in vacuo to afford the title compound as a yellow solid (104 mg, 0.32 mmol, 100%). (*E*)-***S14***: $^1$H NMR (400 MHz, MeOD) δ 7.99 – 7.92 (m, 2H), 7.69 (d, *J* = 2.5 Hz, 1H), 7.66 – 7.60 (m, 2H), 6.64 (d, *J* = 2.5 Hz, 1H), 4.21 (s, 2H), 4.03 (s, 3H). $^{13}$C NMR (151 MHz, MeOD) δ 164.83, 154.36, 137.41, 134.28, 130.99, 124.30, 96.25, 43.92, 39.74. HRMS-EI (m/z) Calculated for $C_{11}H_{14}N_4$ $[M+H]^+$, 216.1244; found 216.1245.

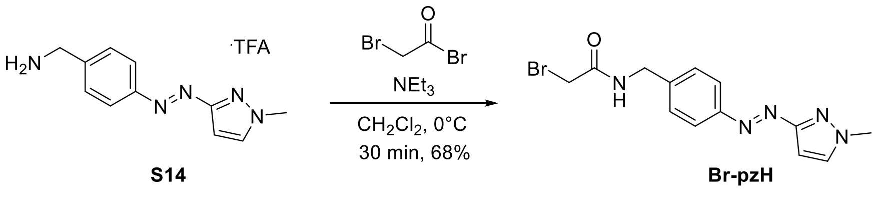

**Br-pzH**. TFA salt **S14** (98 mg, 0.3 mmol) was suspended in $CH_2Cl_2$ (2.5 mL). $NEt_3$ (104 µL, 0.74 mmol, 2.5 eq) was added and the solution cooled to 0 °C and bromo acetylbromide (40 µL, 0.45 mmol, 1.5 eq) was added and the solution was stirred at 0 °C for 30 minutes. The solution was diluted with $CH_2Cl_2$, then the volatiles removed by rotary evaporation (35 °C water bath). The residue was purified by flash-column chromatography (10% EtOAc in $CH_2Cl_2$) to afford the product as a yellow solid (68.2 mg, 202 µmol, 68%). (*E*)-**Br-pzH**: $^1$H NMR (400 MHz, DMSO) δ 8.87 (d, *J* = 6.1 Hz, 1H), 7.83 (d, *J* = 2.4 Hz, 1H), 7.80 (d, *J* = 8.1 Hz, 2H), 7.46 (d, *J* = 8.1 Hz, 2H), 6.53 (d, *J* = 2.4 Hz, 1H), 4.39 (d, *J* = 6.0 Hz, 2H), 3.97 (s, 3H), 3.95 (s, 2H). $^{13}$C NMR (151 MHz, DMSO) δ 166.72, 163.46, 151.56, 143.69, 133.57, 128.64, 122.67, 95.00,

41.83, 40.55, 29.90. HRMS-EI (m/z) Calculated for $C_{13}H_{15}N_5OBr$ $[M+H]^+$, 336.0454; found 336.0457.

##### Synthesis of Br-fAzo

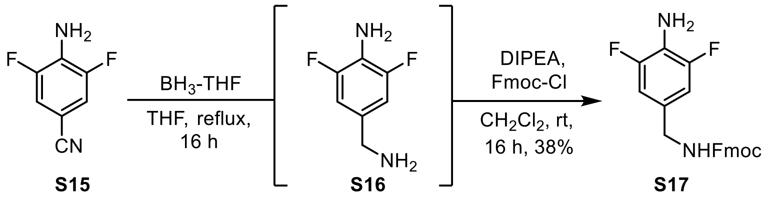

Aniline **S17**. **S15** (1.45 g, 9.41 mmol) was dissolved in THF (20 mL). Borane-THF (1M, 38.4 mL, 38.4 mmol, 4 eq) was added at 0 °C under $N_2$ atmosphere, and the reaction was stirred at reflux temperature 16 hours. MeOH (5 mL) was added dropwise at this temperature until effervescence stopped, and the reaction was stirred at reflux temperature for 30 minutes. The solvent was removed in vacuo, redissolved in EtOAc and washed with water. The organic layer was concentrated. The off-white **S16** was redissolved in $CH_2Cl_2$ (30 mL). DIPEA (1.46 mL, 9.41 mmol, 1 eq) and then Fmoc-Cl (2.43 g, 9.41 mmol, 1 eq) were added and the reaction was stirred for 16 hours at room temperature. The solution was concentrated and purified by silica gel chromatography (6:1 Hexane:Acetone) to afford the title compound as a white solid (1.37 g, 3.60 mmol, 38%). $^1$H NMR (400 MHz, $CDCl_3$) δ 7.76 (d, $J$ = 7.6 Hz, 2H), 7.59 (d, $J$ = 7.5 Hz, 2H), 7.40 (t, $J$ = 7.5 Hz, 2H), 7.31 (t, $J$ = 7.5 Hz, 2H), 6.75 (d, $J$ = 7.4 Hz, 2H), 5.02 (s, 1H), 4.47 (d, $J$ = 6.8 Hz, 2H), 4.30 – 4.01 (m, 3H), 3.70 (s, 2H). $^{13}$C NMR (126 MHz, $CDCl_3$) δ 156.49, 152.05 (dd, $J$ = 242.30 Hz, 8.10 Hz), 143.98, 141.49, 127.86 (t), 127.20, 125.11, 123.24 (t, $J$ = 16.2 Hz), 120.14, 110.28 (dd, $J$ = 15.50 Hz, 6.50 Hz) 66.85, 47.42, 44.28. HRMS-ESI (m/z) Calculated for $C_{22}H_{18}F_2N_2O_2$ $[M+H]^+$, 381.1409; found 381.1408

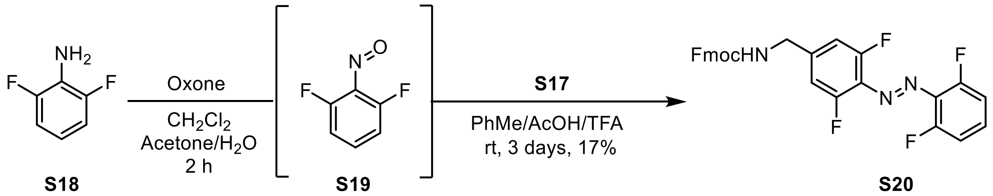

Aniline **S18** (170 mg, 1.31 mmol, 2 eq) was dissolved in $CH_2Cl_2$ (10 mL). Oxone (5.35 g, 13.1 mmol, 20 eq) in water (20 mL) was added and the biphasic solution was vigorously stirred at room temperature for 16 hours, after which the organic layer turned green. The organic layer was separated, then sequentially washed with 1 N HCl (20 mL), sat. bicarbonate solution (20 mL), water (20 mL). The organic layer was dried and concentrated. The residue was redissolved in 6:6:1 AcOH:Toluene:TFA (20 mL), and **S17** (250 mg, 657 µmol, 1 eq) was added, and the solution was stirred at room temperature for 3 days. The mixture was concentrated, then purified by flash-column chromatography (100% $CH_2Cl_2$) to afford the title compound as an orange solid and mixture of isomers (66 mg, 133 µmol, 17%). The sample was irradiated with 405 nm to afford 17:1 ratio of (*E*)-**S20** and (*Z*)-**S20** as indicated by $^1$H NMR analysis. (*E*)-**S20**: $^1$H NMR (600 MHz, $CDCl_3$) δ 7.77 (d, $J$ = 7.1 Hz, 2H), 7.60 (d, $J$ = 7.5 Hz, 2H), 7.41 (t, $J$ = 7.6 Hz, 2H), 7.37 (t, $J$ = 8.9 Hz, 1H), 7.33 (t, $J$ = 7.7 Hz, 2H), 7.06 (t, $J$ = 8.9 Hz, 2H), 6.95 (d, $J$ = 10.4 Hz, 2H), 5.16 (t, $J$ = 6.4 Hz, 1H), 4.55 (d, $J$ = 6.5 Hz, 2H), 4.39 (d, $J$ = 6.4 Hz, 2H), 4.24 (t, $J$ = 6.4 Hz, 1H). $^{13}$C NMR (151 MHz, $CDCl_3$) δ 156.56, 155.89 (dd, $J$ = 263.0, 4.4 Hz), 155.73 (dd, $J$ = 261.4, 4.0 Hz), 143.97, 143.86, 141.55, 132.01, 131.53 (t, $J$ = 10.3 Hz), 130.78, 127.95, 127.25, 125.04, 120.18, 112.77 (dd, $J$ = 20.1, 3.8 Hz), 111.25 (d, $J$ = 20.9 Hz), 66.94 47.48, 44.34. HRMS-EI (m/z) Calculated for $C_{28}H_{20}N_3O_2F_4$ $[M+H]^+$, 506.1486; found 506.1488.

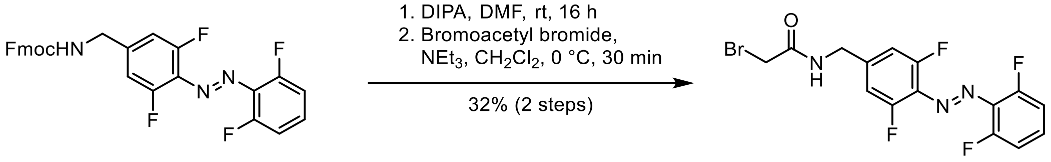

**S20** were dissolved in DMF (66 mg, 131 µmol, 70 mM). DIPA (73 µL, 522 µmol, 4 eq) was added under $N_2$ and the reaction was stirred for 16 hours at room temperature. The reaction was concentrated, then purified by flash-column chromatography to afford the product as an orange solid. The free amine (16 mg, 56 µmol) was dissolved in $CH_2Cl_2$ (40 mM). $NEt_3$ (12 µL, 84 µmol, 1.5 eq) was added and the solution cooled to 0°C and bromoacetylbromide (7 µL, 84 µmol, 1.5 eq) was added and the solution was stirred at 0 °C for 30 minutes. The solution was diluted with $CH_2Cl_2$, then the volatiles removed by rotary evaporation (35 °C water bath). The residue was purified by flash-column chromatography to afford the product as an orange solid (7.3 mg, 18 µmol, 32%). (*E)-**Br-fAzo***: $^{1}H$ NMR (600 MHz, $CDCl_3$) δ 7.37 (tt, *J* = 8.4, 5.8 Hz, 1H), 7.09 – 7.03 (m, 2H), 7.00 (d, *J* = 9.8 Hz, 2H), 6.91 (s, 1H), 4.52 (d, *J* = 6.2 Hz, 2H), 3.98 (s, 2H). $^{13}C$ NMR (151 MHz, $CDCl_3$) δ 165.72, 155.76 (dd, *J* = 262.2, 4.7 Hz), 155.63 (dd, *J* = 261.2, 4.0 Hz), 142.39 (t, *J* = 9.3 Hz), 131.80 (t, *J* = 10.0 Hz), 131.54 (t, *J* = 10.4 Hz), 130.87 (t, *J* = 9.9 Hz), 112.64 (dd, *J* = 20.1, 3.8 Hz), 111.49 (dd, *J* = 21.1, 3.6 Hz), 43.25, 28.88. HRMS-ESI (m/z) Calculated for $C_{15}H_{11}ON_3BrF_4$ $[M+H]^+$, 404.0016; found 404.0017

#### NMR spectra assignment

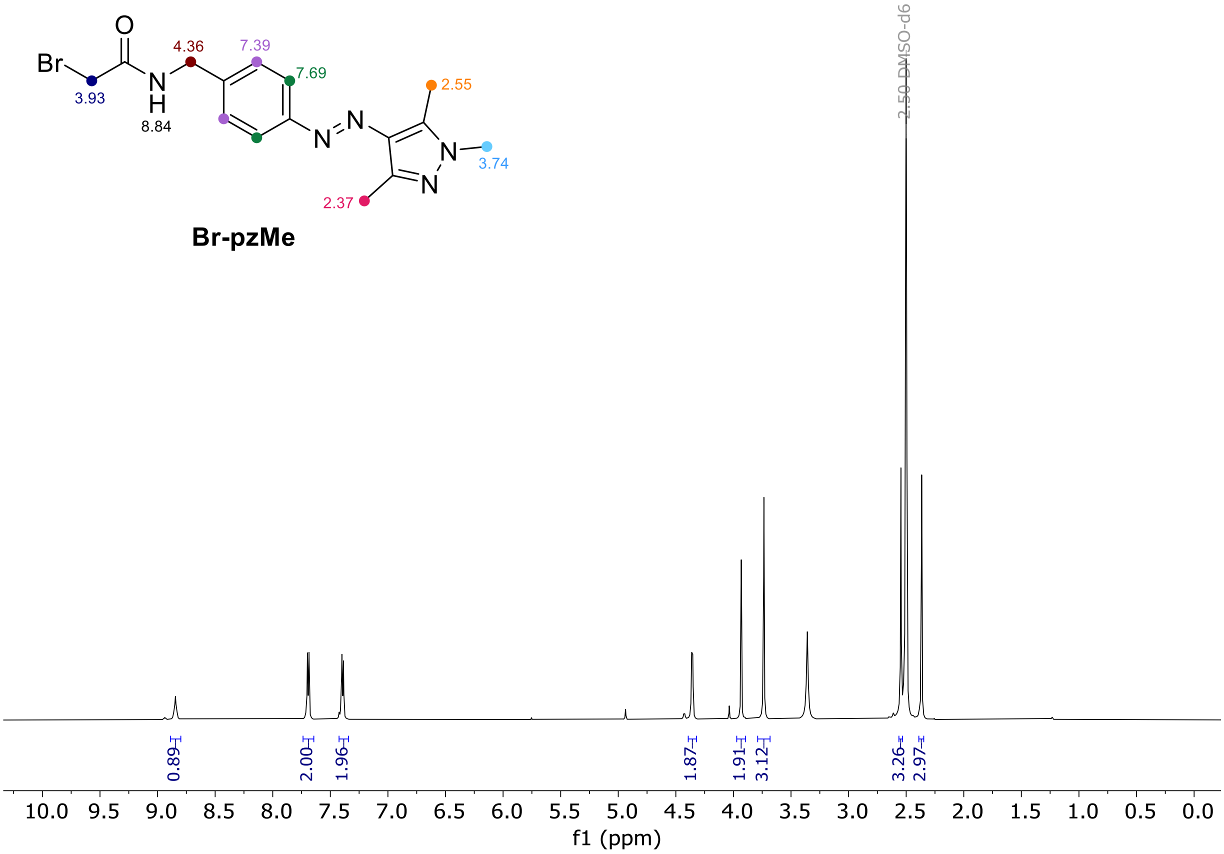

**Fig. 14 |** $^{1}H$ NMR Spectrum of **Br-pzMe** (DMSO-$d_6$, 298 K).

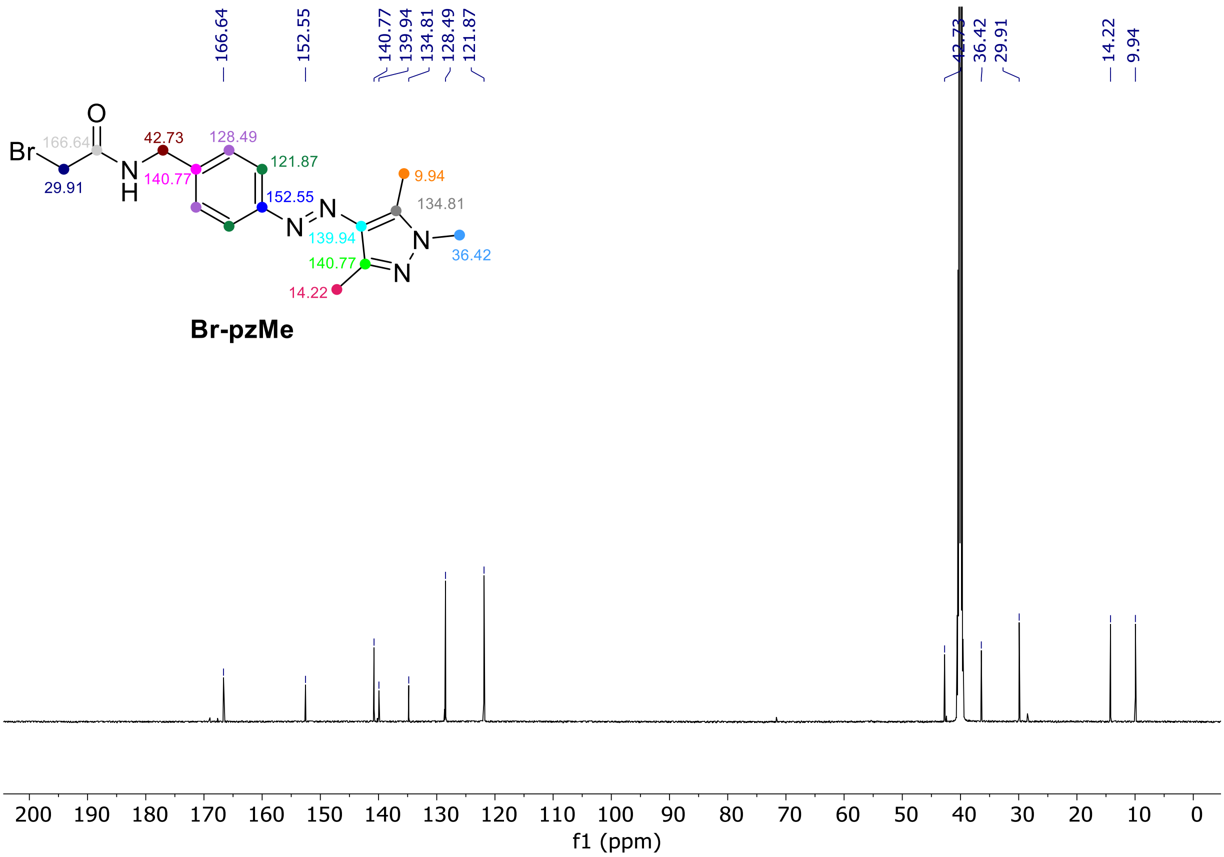

**Fig. 15 |** $^{13}$C NMR Spectrum of **Br-pzMe** (DMSO-$d_6$, 298 K).

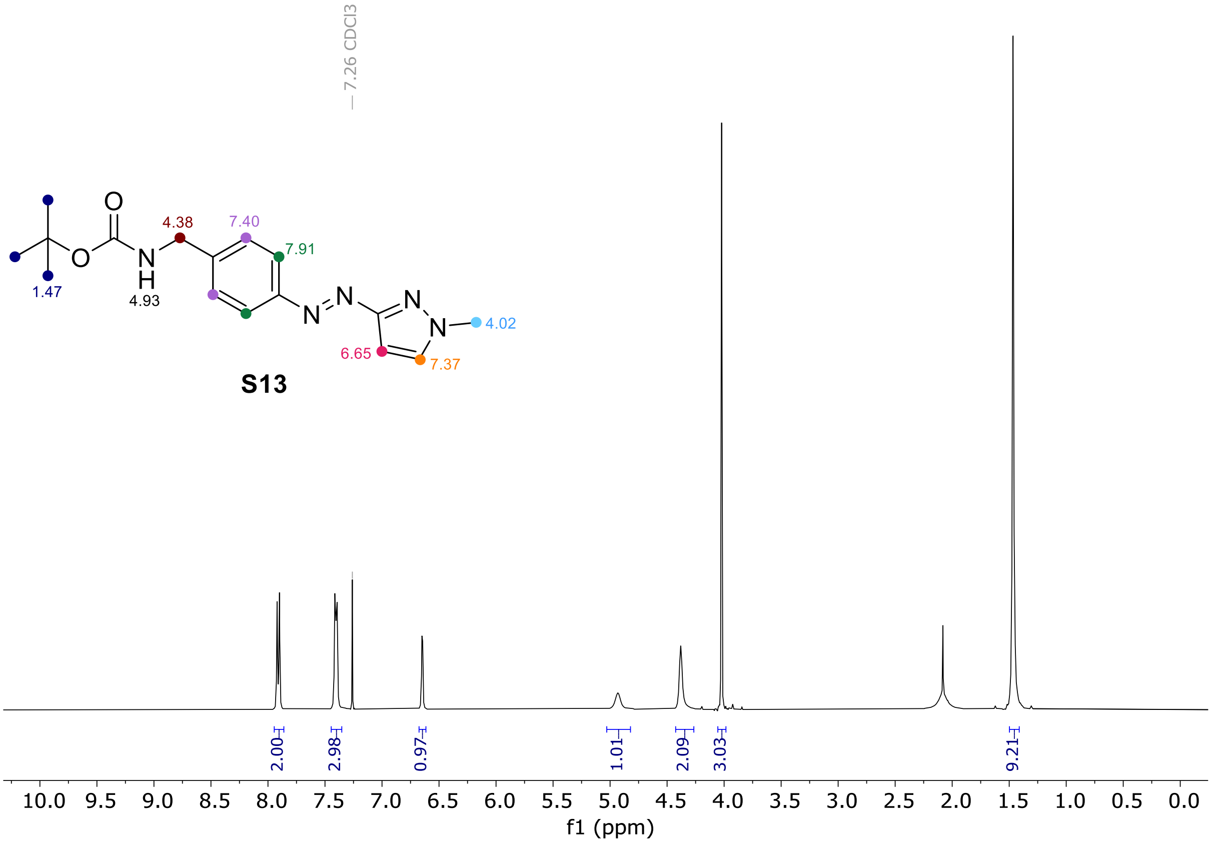

**Fig. 16 |** $^{1}$H NMR Spectrum of **S13** (Chloroform-$d$, 298 K).

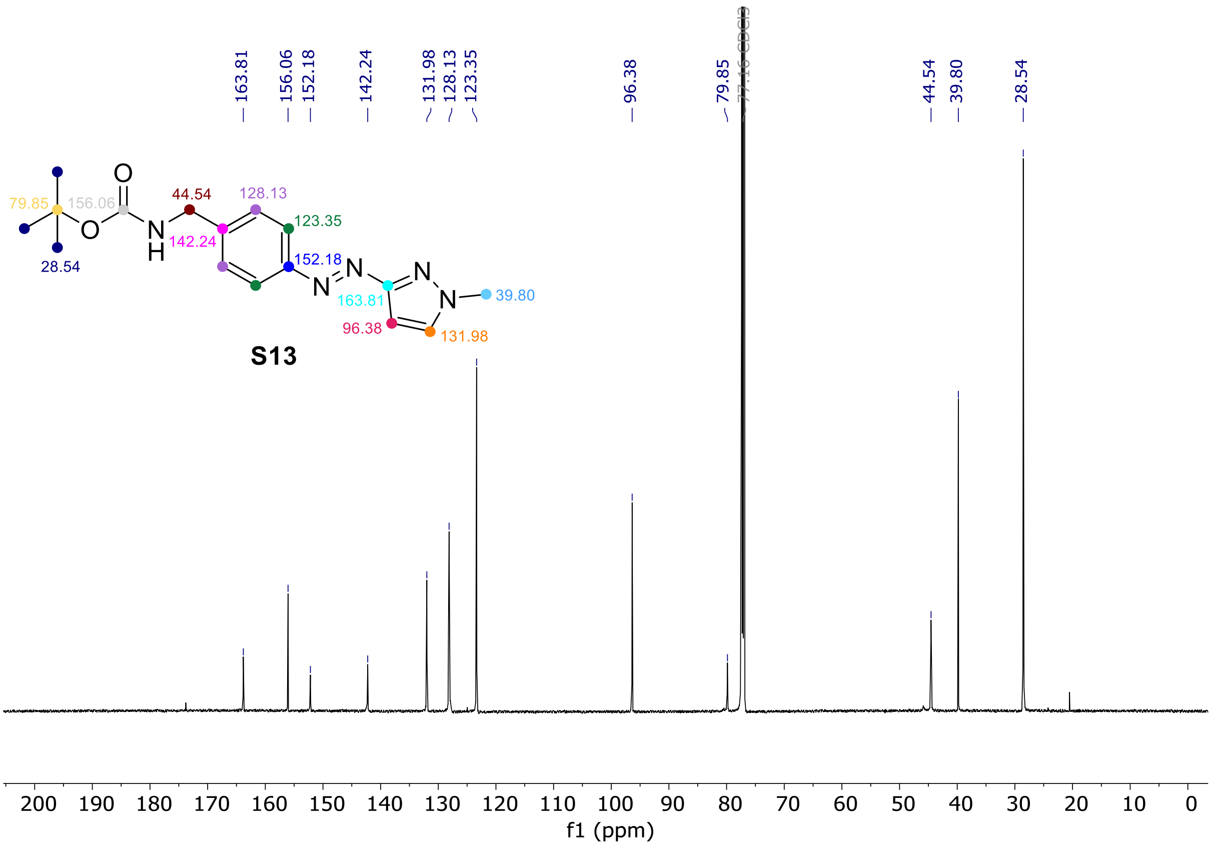

**Fig. 17 |** $^{13}$C NMR Spectrum of **S13** (Chloroform-*d*, 298 K).

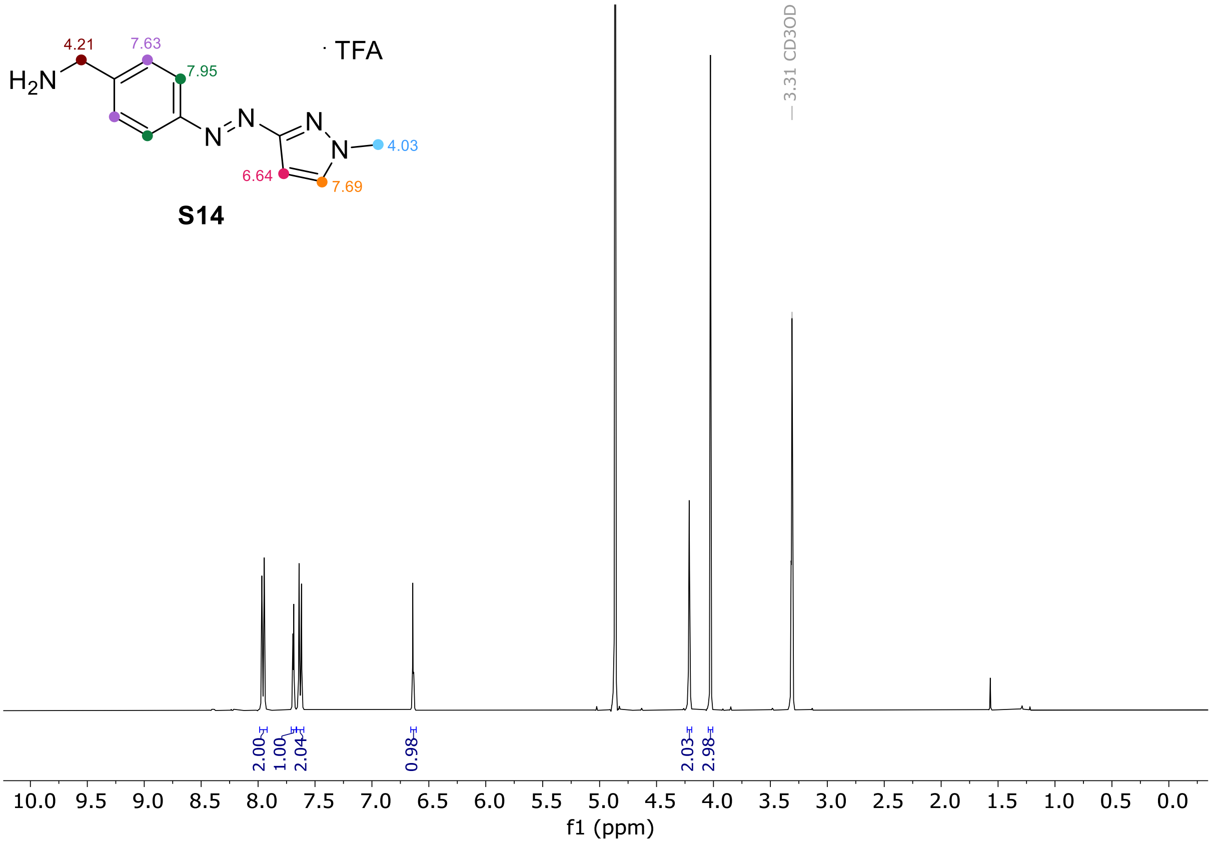

**Fig. 18 |** $^{1}$H NMR Spectrum of **S14** (MeOD-$d_4$, 298 K).

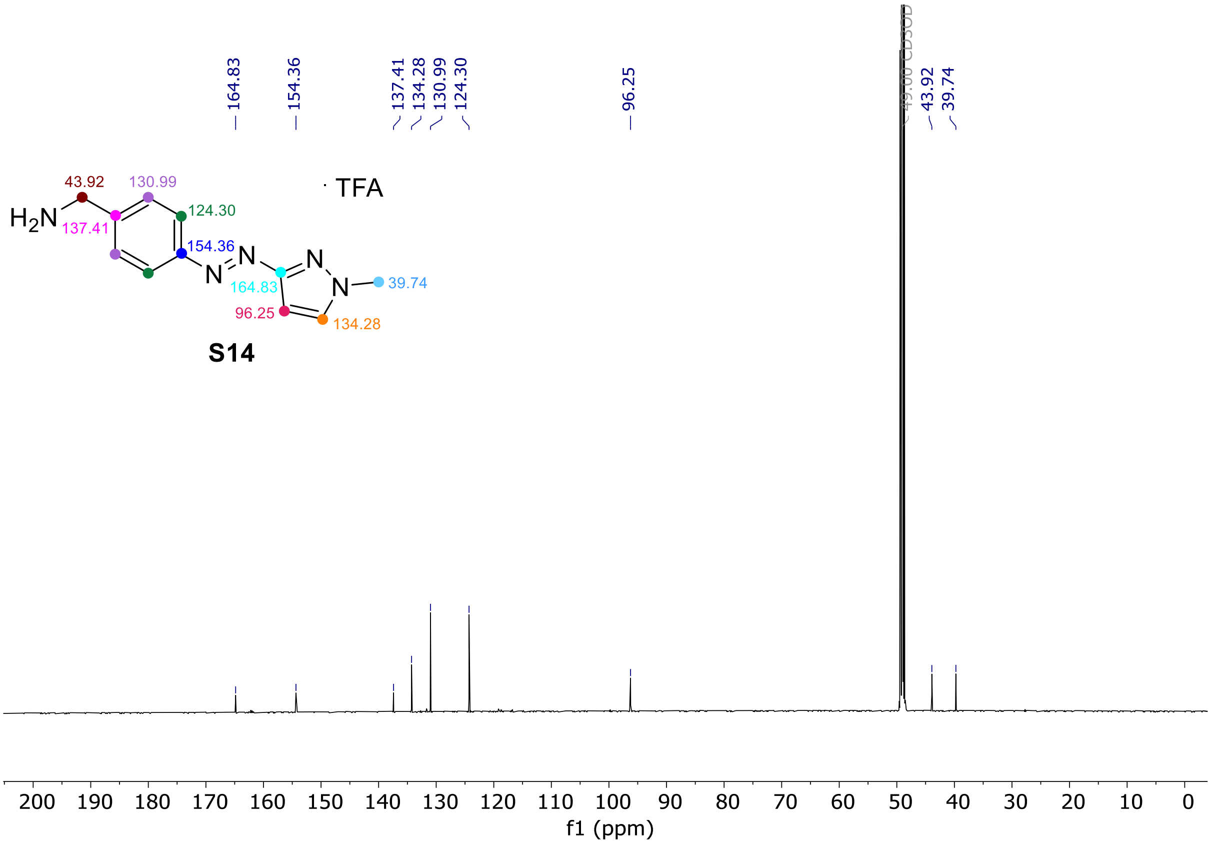

**Fig. 19 |** $^{13}$C NMR Spectrum of **S14** (MeOD-$d_4$, 298 K).

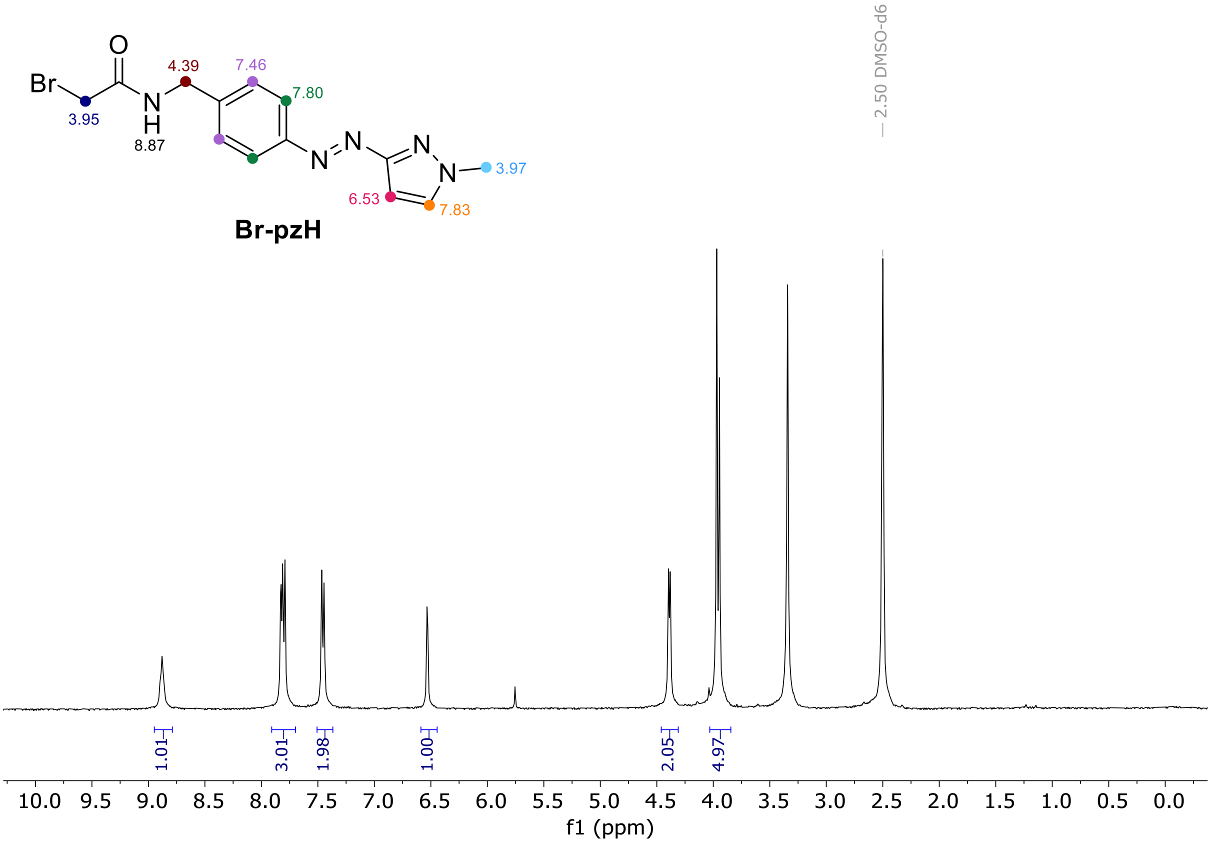

**Fig. 20 |** $^{1}$H NMR Spectrum of **Br-pzH** (DMSO-$d_6$, 298 K).

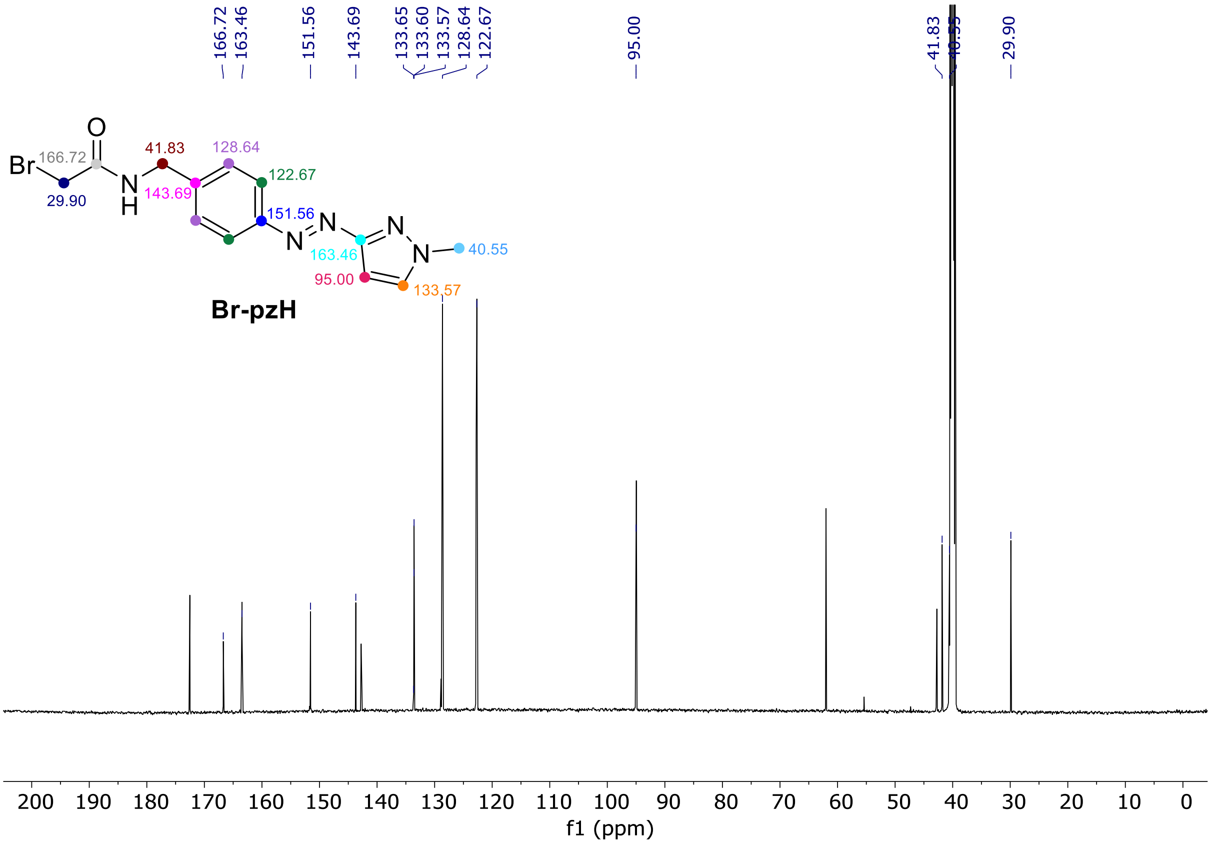

**Fig. 21 |** $^{13}$C NMR Spectrum of **Br-pzH** (DMSO-$d_6$, 298 K).

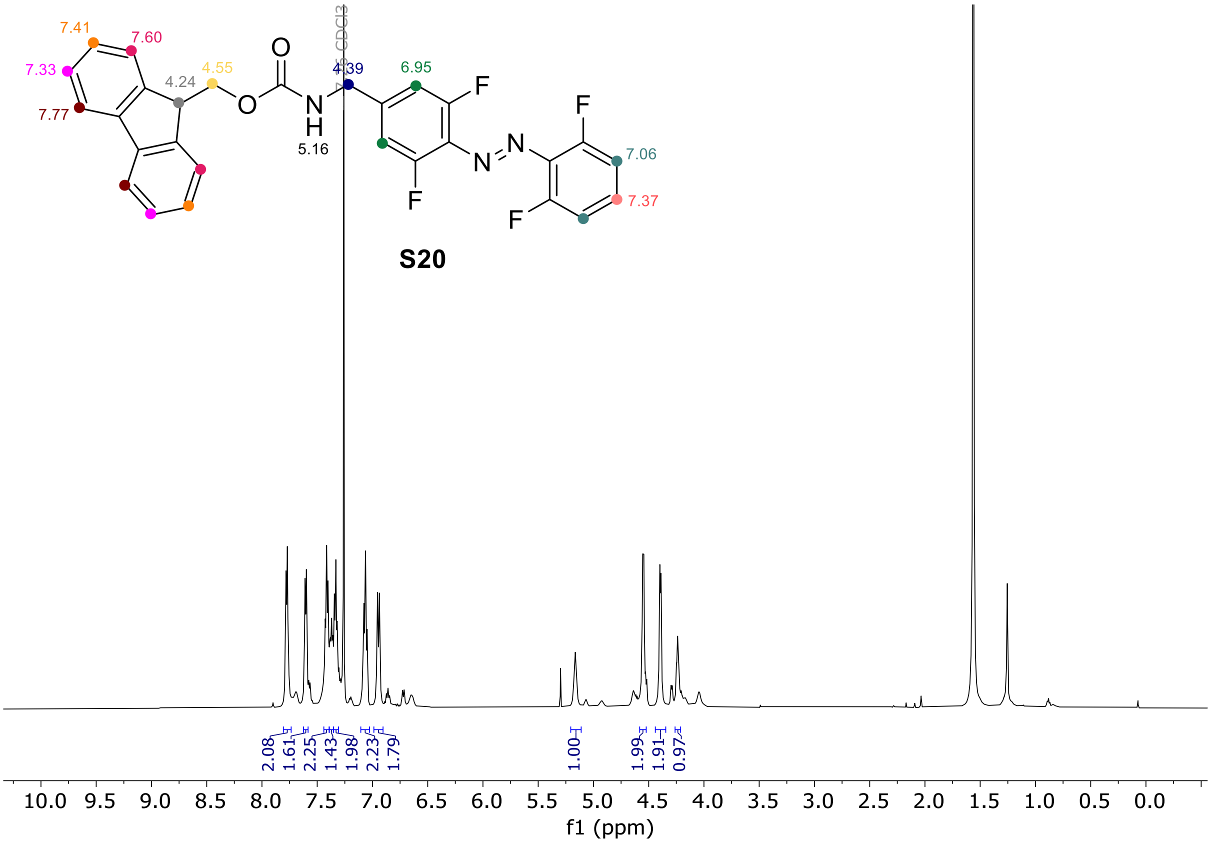

**Fig. 22 |** $^1$H NMR spectrum of **S20.** (Chloroform-$d$, 298 K).

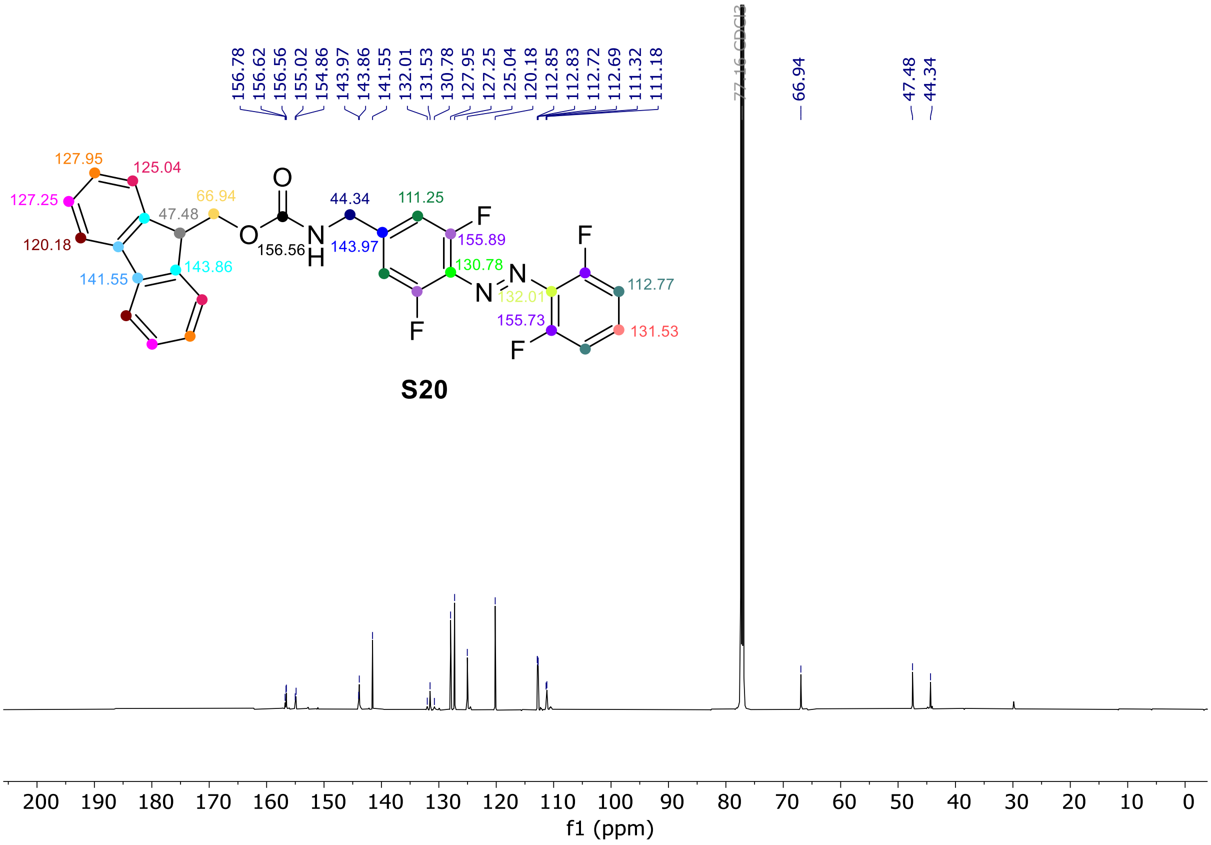

**Fig. 23 |** $^{13}$C NMR spectrum of **S20.** (Chloroform-*d*, 298 K).

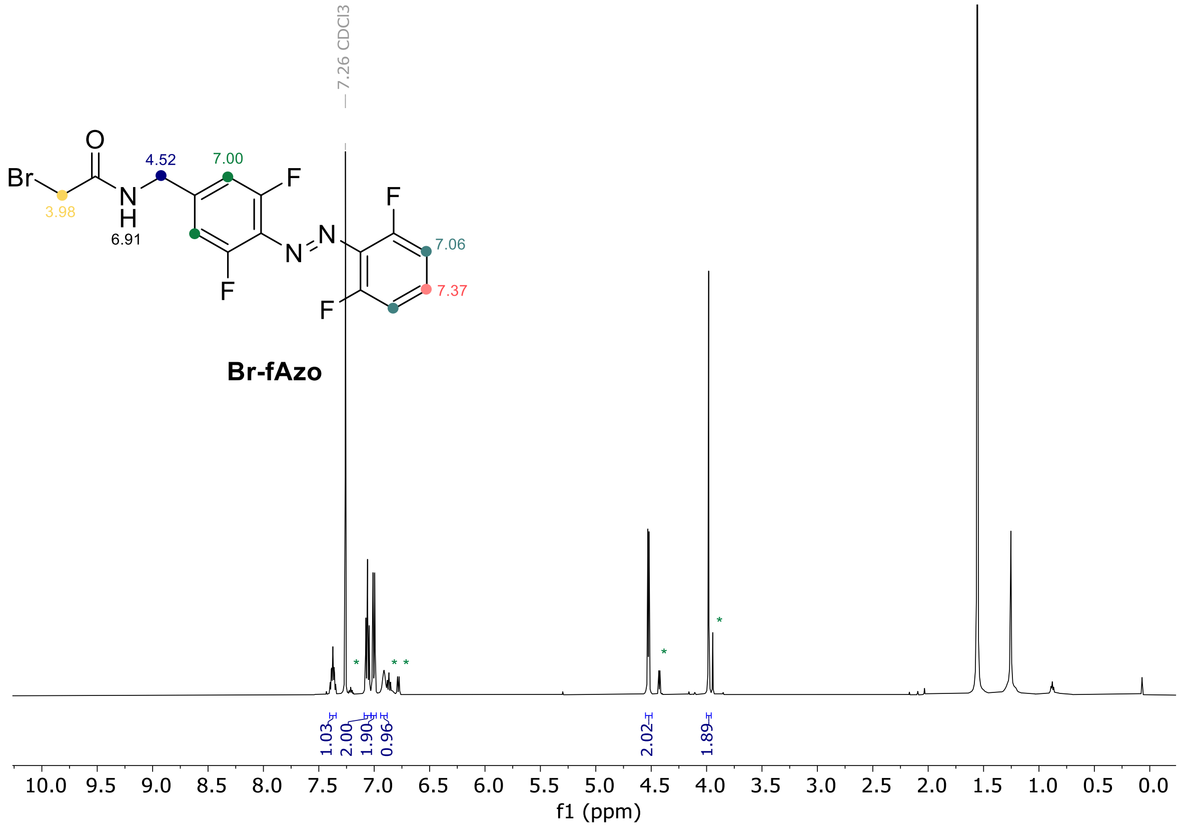

**Fig. 24 |** $^{1}$H NMR spectrum of **Br-fAzo.** (*Z*)-**Br-fAzo** signals labelled as *. (Chloroform-*d*, 298 K).

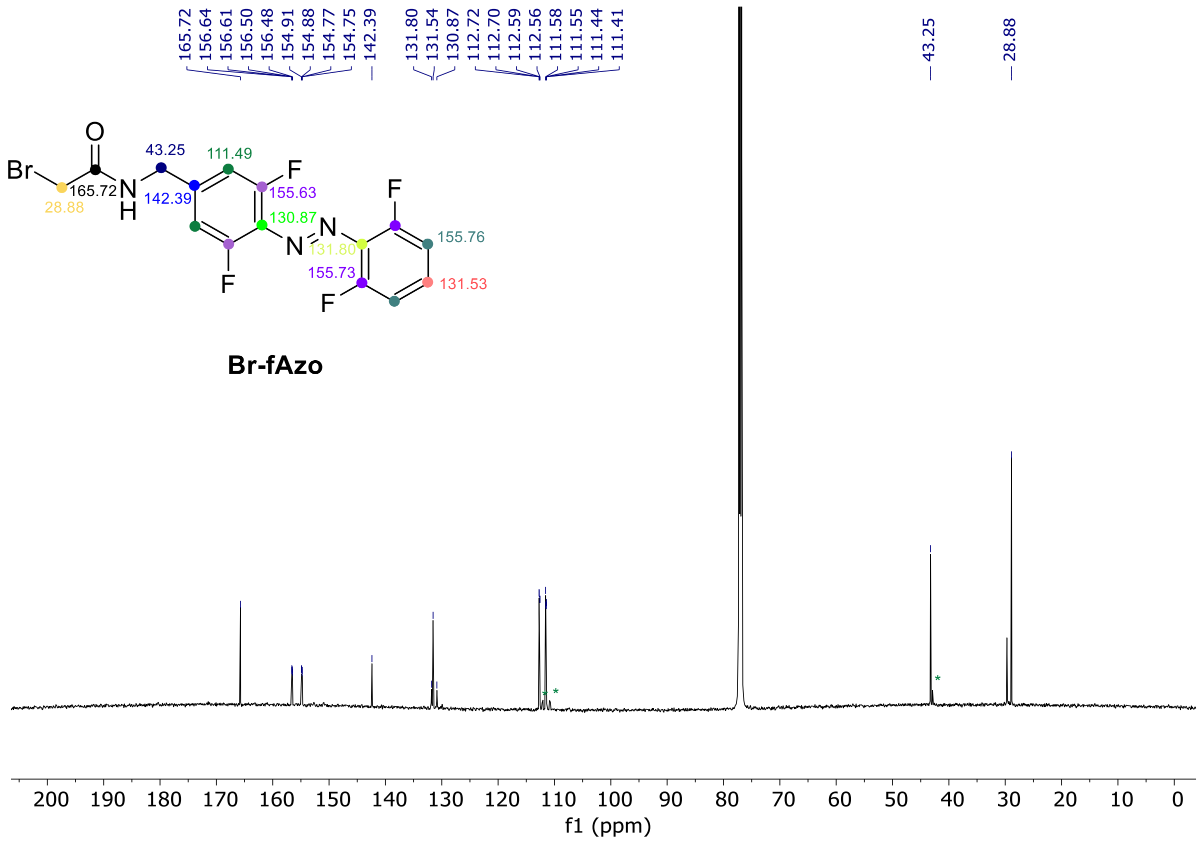

**Fig. 25 |** [13]C NMR spectrum of **Br-fAzo.** (*Z*)-**Br-fAzo** signals labelled as *. (Chloroform-*d*, 298 K).

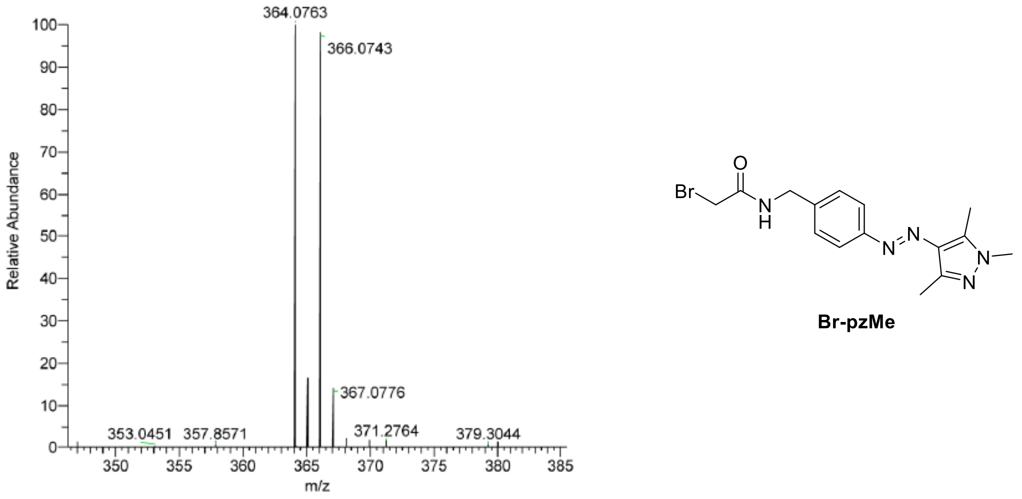

**Fig. 26 |** HRMS spectrum of **Br-pzMe**. HRMS-EI (m/z) Calculated for $C_{15}H_{19}N_5OBr$ $[M+H]^+$ , 364.0767; found 364.0763.

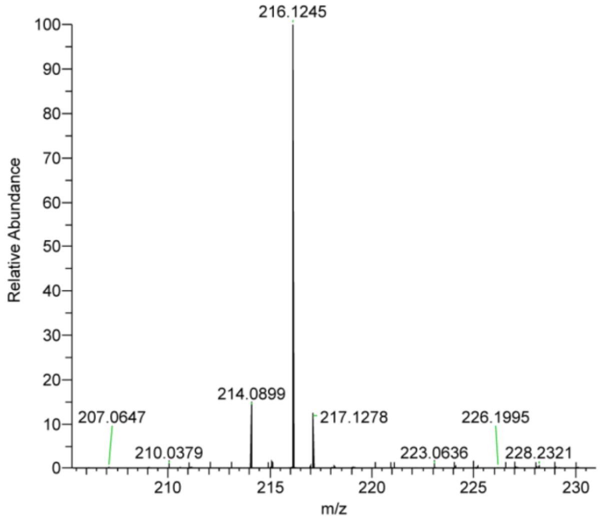

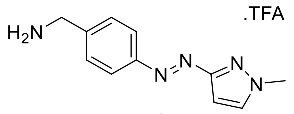

**S14**

**Fig. 27 |** HRMS spectrum of **S14**. HRMS-EI (m/z) Calculated for $C_{11}H_{14}N_4$ $[M+H]^+$, 216.1244; found 216.1245.

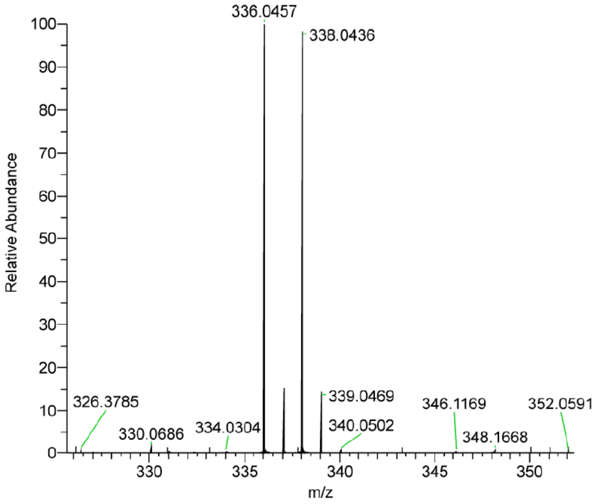

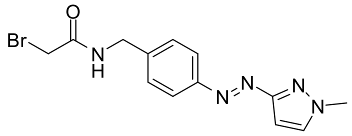

**Br-pzH**

**Fig. 28 |** HRMS spectrum of **Br-pzH**. HRMS-EI (m/z) Calculated for $C_{13}H_{15}N_5OBr$ $[M+H]^+$, 336.0454; found 336.0457.

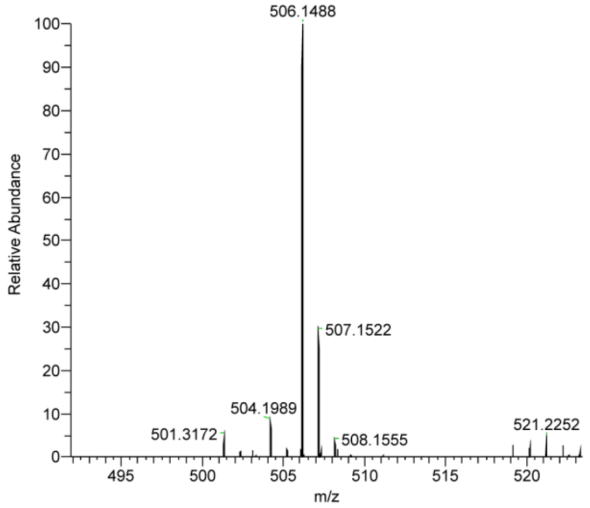

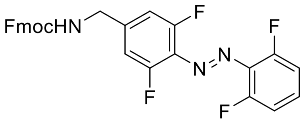

**S20**

**Fig. 29 |** HRMS spectrum of **S20**. HRMS-EI (m/z) Calculated for $C_{28}H_{20}N_3O_2F_4$ $[M+H]^+$, 506.1486; found 506.1488.

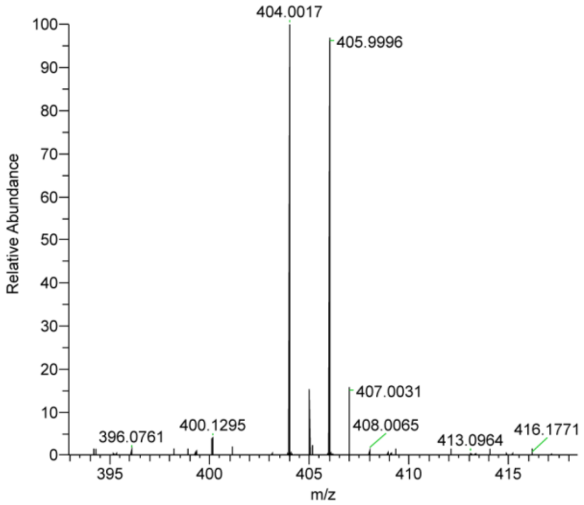

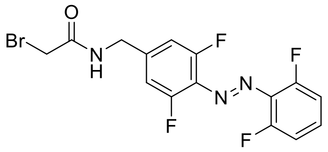

**Br-fAzo**

**Fig. 30 |** HRMS spectrum of **Br-fAzo**. HRMS-ESI (m/z) Calculated for $C_{15}H_{11}ON_3BrF_4$ $[M+H]^+$, 404.0016; found 404.0017
